# Mechanisms underlying the loss of migratory behaviour in a long-lived bird

**DOI:** 10.1101/2024.03.06.583673

**Authors:** Pedro Andrade, Aldina M. A. Franco, Marta Acácio, Sandra Afonso, Cristiana I. Marques, Francisco Moreira, Miguel Carneiro, Inês Catry

**Affiliations:** CIBIO, Centro de Investigação em Biodiversidade e Recursos Genéticos, InBIO Laboratório Associado, Campus de Vairão, Universidade do Porto, 4485-661 Vairão, Portugal; BIOPOLIS Program in Genomics, Biodiversity and Land Planning, CIBIO, Campus de Vairão, 4485-661 Vairão, Portugal; School of Environmental Sciences, University of East Anglia, NR4 7TJ, Norwich, Norfolk, UK; School of Zoology, Faculty of Life Sciences, Tel Aviv University, Tel Aviv 69978, Israel; Departamento de Biologia, Faculdade de Ciências, Universidade do Porto, 4099-002 Porto, Portugal; CIBIO, Centro de Investigação em Biodiversidade e Recursos Genéticos, InBIO Laboratório Associado, Instituto Superior de Agronomia, Universidade de Lisboa, 1349-017 Lisbon, Portugal

**Keywords:** bird migration, GPS-tracking, adaptation, phenotypic flexibility, developmental plasticity, ontogeny, generational shifts, white stork

## Abstract

Human-induced environmental changes are shifting the migration patterns of birds worldwide. Species are adjusting migration timings, shortening and diversifying migratory routes, or even completely disrupting migration and transitioning towards residency. Whilst the ultimate causes driving changes in migratory patterns are well established, the underlying mechanisms by which migratory species adapt to environmental change remain unclear.

Here, we studied the mechanisms driving the recent and rapid loss of migratory behaviour in Iberian white storks *Ciconia ciconia*, a long-lived and previously fully migratory species through the African-Eurasian flyway. We combined 25 years of census data, GPS-tracking data from 213 individuals (80 adults and 133 first-year juveniles) followed for multiple years, and whole-genome sequencing, to disentangle whether within- (phenotypic flexibility) or between- (developmental plasticity or microevolution, through selection) individual shifts in migratory behaviour over time can explain the observed population-level changes towards residency.

Between 1995 and 2020, the proportion of individuals no longer migrating and remaining in Southern Europe year-round increased dramatically, from 18% to 68-83%. We demonstrate that this behavioural shift is likely explained by developmental plasticity. Within first-year birds, 98% crossed the Strait of Gibraltar towards their African wintering grounds, in Morocco or Sub-Saharan Africa. However, the majority shifted towards a non-migratory strategy as they aged - the proportion of migrants decreased to 67% and 33%, on their second and third year of life, respectively - suggesting that migratory behaviour is determined during ontogeny. Supporting these findings, only 19% of GPS-tracked adults migrated. Moreover, we did not find evidence of phenotypic flexibility, as adults were highly consistent in migratory behaviour over multiple years (only 3 individuals changed strategy between years, out of 113 yearly transitions), nor of selection acting on genetic variation, since genomes of migrants and residents are essentially undifferentiated.

Our results suggest that through developmental plasticity, traits that are plastic during specific windows of development, become fixed during adulthood. Thus, inter-generational shifts in the frequency of migratory and non-migratory young individuals could drive population changes in migratory behaviour. This can provide a fast mechanism for long-lived migratory birds to respond to rapid human-driven environmental changes.

## 1. INTRODUCTION

The migratory behaviour of many bird species is shifting in response to human-induced environmental change. This includes changes in migratory phenology, in the limits of breeding and wintering ranges, the shortening of migration distances or even the disruption of migration (Able & Belthoff, 1998; Berthold et al., 1992; Curley et al., 2020; Horton et al., 2020; Rushing et al., 2020; Visser et al., 2009). While the ultimate causes driving observed changes in migratory patterns are well established (e.g. human-driven climate change), the mechanisms by which migratory species adapt to environmental change remain largely unknown (Åkesson &. Helm, 2020; Charmantier & Gienapp, 2014). At the population level, changes in avian migratory behaviour can occur through three distinct but not mutually exclusive processes: (i) phenotypic flexibility, induced by environmental conditions, which can be reversible and reflects within-individual changes in the expression of a phenotype (Charmantier et al., 2008); (ii) evolution, through longer-term, inter-generational shifts in the frequency of alleles controlling migratory behaviour (Berthold et al., 1992); and (iii) developmental plasticity, an irreversible phenotypic change induced by environmental, physiological, or social conditions juveniles experience during ontogeny (Piersma & Drent, 2003). Changes at the population level are then driven by generational shifts, with young individuals replacing old ones in the population.

Cross-breeding experiments in passerines, conducted in captivity, indicate substantial heritability of migratory traits (Pulido et al., 2001, 2010). Heritable differences in migratory strategies have been tentatively linked to either large genomic regions of high differentiation, likely harbouring chromosomal inversions containing dozens to hundreds of genes (Delmore et al., 2016; Lundberg et al., 2017; Sanchez-Donoso et al., 2022; Sokolovskis et al., 2023), or to selection operating on genes that are potentially linked to spatial behaviour and learning (Delmore et al., 2020; Gu et al., 2021; Toews et al., 2019). Although these examples help to illustrate how genetic processes may lead to changes in migratory behaviour through genetics, plasticity could provide a faster mechanism for rapid acclimation to environmental change (Åkesson &. Helm, 2020; Both & Visser, 2001; Charmantier et al., 2008; Charmantier & Gienapp, 2014; Dias et al., 2011; Horton et al., 2023; Teitelbaum et al., 2016; Teplitsky et al., 2008), which is particularly relevant for long-lived species since important environmental changes can occur within the lifetime of individuals.

Attempts to disentangle the relative roles of different modes of plasticity in the evolution of bird migration are hampered by the difficulty of collecting long-term population-wide information, as well as repeated individual data through periods of shifts in migratory behaviour at the population level (Conklin et al., 2021; Fraser et al., 2019; Gill et al., 2019). Recently, developments in tracking technology warranted to bridge this knowledge gap are becoming more widely available (Åkesson &. Helm, 2020). However, recent research using repeat-tracking data does not unambiguously support a single mechanism. Some studies have shown that phenotypic flexibility can largely account for population-level advances in migration timing of long-distance migrants (e.g., Conklin et al., 2021; Gill et al., 2019). Yet, a growing number of studies document high within-individual levels of repeatability in migratory timing, routes, and wintering sites over multiple years (Carneiro et al., 2019; Pederson et al., 2018; Stanley et al., 2012; Vardanis et al., 2011), with some of them arguing against heritable variation as the causal mechanism. This suggests that, for species with low individual flexibility, generational shifts through development plasticity could drive population-level changes in migratory routes and timing in few generations, with young individuals being the agents of such change (Gill et al., 2019; Verhoeven et al., 2018, 2021). These studies, however, have not explicitly discounted the effects of underlying genetic variation. Long-lived birds, in particular, have extended developmental periods in which young birds can learn from past experiences and improve their migratory performance (Campioni et al., 2020; Sergio et al., 2014). Moreover, in social species, individuals can alter their migratory behaviour based on social learning from conspecifics, also contributing to population-level shifts in migratory behaviour (Mueller & O’Hara, 2013; Teitelbaum et al., 2016).

Many studies investigating recent changes in avian migratory patterns have focused mostly on shifts in routes or timing of migration (time and space). However, even more profound changes have been recorded, with individuals abandoning migration completely in previously partially or fully migratory populations and establishing non-migratory populations (Berthold, 2001; Newton, 2008; Van Vliet et al., 2009). Whilst these changes can dramatically affect species distributions and ecosystems with unforeseen conservation challenges (Gill et al., 2019), the mechanisms underlying the rapid losses of migratory behaviour are poorly understood. Here, we combine two decades of census data, movement information from 213 birds GPS-tracked for up to 7 years and whole-genome sequencing to investigate the loss of migratory behaviour in the white stork (*Ciconia ciconia*, Figure 1A). This long-lived species is an iconic symbol for long-distance migrations in many European countries, where juvenile and adults migrate together in mixed flocks (Rotics et al., 2018), as naïve birds require guidance from older and more experienced individuals to successfully reach their wintering grounds (Dallinga & Schoenmakers, 1987). In recent decades, however, a growing number of individuals no longer carry out their annual autumn migration from Europe to Africa, remaining in Southern Europe all year-round (Archaux et al., 2004; Cheng et al., 2019; Flack et al., 2016; Rotics et al., 2017). Increased food availability from anthropogenic sources, including food waste at landfills (Gilbert et al., 2016; Soriano-Redondo et al., 2021; Tortosa et al., 2002) and the invasive red crayfish *Procambarus clarkii* (Ferreira et al., 2019), together with increases in winter temperatures (Schulz & Schulz, 1999) are thought to have contributed to the suppression or shortening of migration in European white storks. Nonetheless, the mechanisms underlying this behavioural shift at the population-level remain unknown.

**Figure 1.**
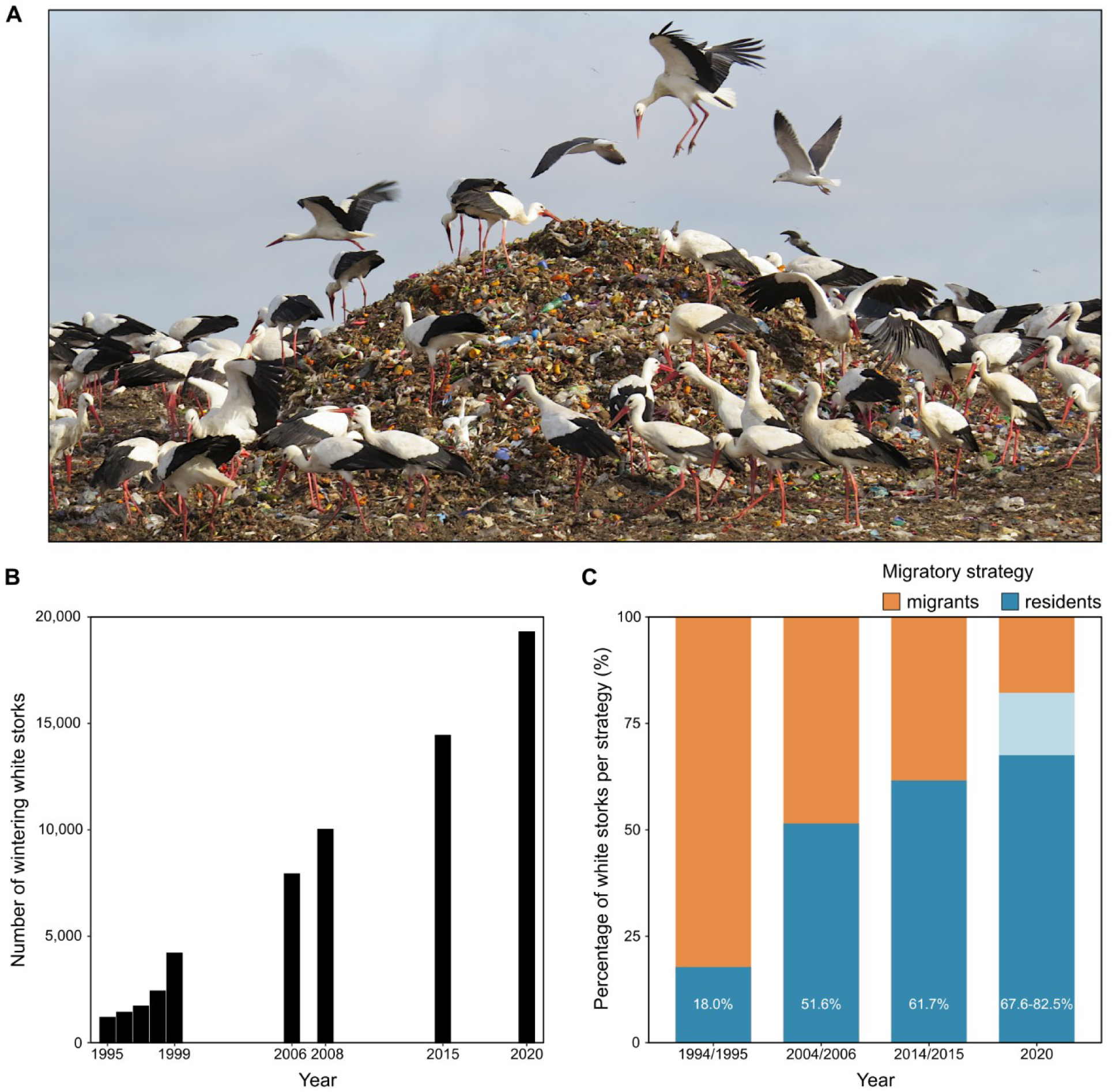
Loss of migratory behaviour in the white stork *Ciconia ciconia*. (A) Increased food availability, particularly at landfills, is one of the major factors promoting an increase in the wintering population of white storks in Europe (Catry et al., 2017; Cheng et al., 2019; Flack et al., 2016). (B) Population trends of wintering white storks from the Portuguese population between 1995-2020 (Catry et al., 2017), and this study). (C) Changes in the percentage of resident white storks in Portugal in the last 25 years. The proportion of residents was estimated as the ratio between the number of breeding and wintering individuals (see “Methods”). To estimate the proportion of residents in 2020 (no breeding census was performed since 2014) we used the number of breeding pairs in 2014 (minimum value) and the predicted number of breeding pairs given by the linear model (y = 838.9x + 5890.1) using the data available from the previous census (maximum value). The range of this estimate is represented by a lighter shade of blue.

National surveys performed during the last 25 years (Catry et al., 2017) and additional data from the present study) show that the number of Portuguese white storks remaining in the country during the winter increased 16-fold from 1,187 individuals in 1995 to 19,295 in 2020 (Figure 1B), coinciding with a northward wintering area range expansion (Figure S1). The breeding population increased 3.5 times in the same period (from 3,302 in 1994 to 11,691 breeding pairs in 2014; (Encarnação, 2015), indicating a steep rise in the proportion of resident (i.e., non-migratory) individuals, from 18% to 68-83% (Figure 1C, Figure S2). Here, we aim to understand the mechanisms driving the observed population-level changes in migratory behaviour of white storks. Specifically, we investigate whether such changes can derive from: (i) within-individual reversible changes in the expression of the phenotype (phenotypic flexibility); (ii) between-individual, non-reversible changes in the expression of the phenotype, linked to developmental effects (developmental plasticity); and (iii) selection on genetic variation (microevolution).

## 2. MATERIALS AND METHODS

### 2.1. Population trends of resident white storks

White stork national censuses have been carried out in Portugal over the last 25 years, to monitor the species population trends following declines that occurred earlier during the 20^th^ century. Breeding censuses were performed in 1994, 2004 and 2014 by quantifying the number of occupied nests in spring (Encarnação, 2015). Non-breeding censuses were conducted from mid-September to early October from 1995-1999 (annually) and in 2006, 2008, 2015 (Catry et al., 2017) and 2020 (this study) and included all areas where the species is known to winter regularly, particularly areas of high food availability during winter, such as landfill sites and rice fields, where storks tend to concentrate (Catry et al., 2017). The census period was chosen because most migratory individuals cross the Strait of Gibraltar towards their African wintering grounds between July and early September (Acácio et al., 2022; Fernández-Cruz, 2004; Soriano-Redondo, 2020), and the pre-nuptial return migration to the breeding areas only starts in November (Bécares et al., 2019, authors’ tracking data). Moreover, from mid-September to mid-October, the number of immigrants is very small, as suggested by tracking data and ring resights of non-Portuguese individuals (Supplementary text, Figure S2). Thus, all birds counted during the non-breeding surveys were considered as residents (see Supplementary text for a more detailed description). For further validation, we estimated the proportion of resident birds using data from our GPS-tracked adult population (see below) between 2016 and 2022 and compared it with the estimates from the national surveys.

### 2.2. GPS tracking of white storks

We captured 213 white storks (80 adults and 133 juveniles) in southern Portugal between 2016 and 2022 to deploy GPS-tracking devices. Storks were tracked for a minimum of 2 months and up to 7 years, enough to identify at least one migratory strategy per individual. Adult white storks were captured either at their nests, using remotely activated clap nets, or at landfill sites using nylon leg nooses. Juveniles were taken from the nests for tag deployment and returned afterwards. All captured individuals were fitted with individually coded rings, measured, and blood samples were collected and stored in ethanol. GPS/GSM loggers (“Flyway-50” from Movetech Telemetry, “Ornitrack-50’ from Ornitela and 50g bird solar tags from e-obs GmbH) were mounted on the back of the birds as backpacks with a Teflon harness; the total weight was 50-90 g, 1.5–3.7% of the birds’ body mass. The loggers were programmed to record GPS positions every 20 min. The procedure was approved by Instituto da Conservação da Natureza e das Florestas (Portugal, permits 493/2016/CAPT, 662/2017/CAPT, 549/2018/CAPT, 248/2019/CAPT, 365/2020/CAPT, 199/2021/CAPT and 542/2022/CAPT). Migrants were defined as birds that crossed the Strait of Gibraltar (to Morocco or Sub-Saharan Africa), while residents remained in Iberia all year-round (Portugal and Spain).

### 2.3. Ontogeny and consistency in migratory behaviour of white storks

To investigate shifts in migratory decisions of white storks with age, we fitted a binomial generalised linear mixed model (g*lmer* function in the *R-*package *lme4,* (Bates et al., 2014) with migration strategy (resident or migrant) and age (*n* = 133, 24, 12 and 83 individuals with 1, 2, 3 and ≥4 years) as the response and explanatory variables, respectively and bird ID as random effect. Birds tagged as breeding adults that changed migratory strategy were not included in the analysis (n=3).

To assess the degree of individual consistency (or conversely, flexibility) in the choice of wintering grounds (and thus, on migratory strategy) we used data from adults and juveniles tracked in multiple years. For each bird, we determined the wintering latitude as the minimum latitude reached in October, as by then white storks are in their main wintering grounds and do not show significant dispersive or migratory movements (Rosa et al., 1998). Amongst adults, 48 individuals were tracked for at least two consecutive winters; 11, 17, 15, 3, 1 and 1 storks were repeatedly tracked through 2-7 winters, respectively (161 bird-winter comparisons). Within juveniles, we tracked 24 juveniles for at least two consecutive winters; 13, 5, 3, 1, 1, 1 storks were repeatedly tracked for 2-7 winters, respectively. However, to estimate individual repeatability amongst juveniles, we discarded bird-winters after the age of first breeding (remaining with 59 bird-winter comparisons). Repeatability of wintering latitudes was analysed using the *rpt* function for Gaussian data in the *R*-package *rptR* (Stoffel et al., 2017).

### 2.4. Reference genome assembly

We assembled the genome of the white stork using linked read technology (10X Genomics, San Francisco, USA). Snap-frozen fresh blood from a female bird (metal ring number MR09149, CEMPA), over-wintering within the Iberian Peninsula, was used for high-molecular weight DNA extraction using a salt-based protocol (Enbody et al., 2021). Prior to library preparation, DNA quantity and integrity was assessed using a NanoDrop instrument, Qubit dsDNA BR Assay Kit and Agilent Genomic DNA ScreenTape (Agilent). The average size of the extracted DNA was estimated to be above 60 kb. A Chromium library was prepared by Novogene UK following manufacturer’s instructions, and sequenced on a Illumina instrument using 2×150 bp paired-end reads. This produced a total of 510,084,146 reads (corresponding to an effective coverage of 60.9X). Chromium sequencing data are available in the Sequence Read Archive under BioProject PRJNA713582.

To assemble the genome using linked-read data, we used *Supernova* v2.0 (Weisenfeld et al., 2017) with default parameters. A preliminary version of the assembly was subjected to an automated Contamination Screen from NCBI’s genome submission portal identified a number of sequences that were either duplicated, of foreign origin or mitochondrial, and these were removed from the assembly. Adaptor sequences (NGB01088.1, NGB01096.1, NGB01047.1 and NGB01039.1) were also identified flanking gaps in 20 independent regions of the assembly, suggestive of local misassemblies. To test for this, we mapped the whole genome re-sequencing dataset from 54 birds (see below) to the preliminary assembly and used *IGV* v2.8.2 (Thorvaldsdóttir et al., 2013) to look for pairs of reads spanning each gap. 19 out of 20 of these regions were considered to result from correct assembly, since for each of them we found at least 10 pairs of reads that respected the following criteria: 1) each read mapped on a different side of the gap; 2) pairs of reads had mapping quality 60; and 3) reads had no supplementary or secondary alignments. Following this inspection, adaptor sequences were hard-masked. For the remaining region a large gap meant it was not possible to rule out a misassembly, so we split the scaffold into two smaller ones.

To evaluate the quality of our final assembly, we calculated summary metrics with the script *assemblathon_stats.pl* (https://github.com/KorfLab/Assemblathon (Bradnam et al., 2013). We also quantified the number of highly conserved single-copy orthologues using *BUSCO* v5 (Simão et al., 2015). For *BUSCO* analysis, we ran the *MetaEuk* gene predictor (Levy et al., 2020), with the lineage dataset aves_odb10. To identify coordinates for protein coding sequences we downloaded the reference assembly and annotation of the ruff (*Calidris pugnax*) obtained from NCBI (GCF_001431845.1) and conducted a lift-over of the coordinates from the annotation GFF3 file through *Liftoff* v1.6.3 (Shumate & Salzberg, 2021), with options -polish and -flank 0.1, using *minimap2* v2.17-r941 (Li et al., 2018) to align gene sequences.

### 2.5. Whole-genome re-sequencing, read mapping and variant calling

We re-sequenced the genomes of multiple individuals at low-coverage using short-read high-throughput sequencing. Based on the data we obtained through GPS tracking, we selected 54 adult individuals (11 migrants, 43 residents). Genomic DNA was extracted from blood stored in ethanol (using the same salt-based extraction protocol as mentioned above), and DNA quantity and integrity was assessed using a NanoDrop instrument, Qubit dsDNA BR Assay Kit and through agarose gel visualization. Individual whole-genome libraries were prepared using the TruSeq DNA PCR-Free kit (Illumina). Libraries were quantified by qPCR using the KAPA Library Quantification Kit, pooled, and sequenced to an average depth of 2.1X using 2×150 bp reads in a Illumina instrument at Novogene UK. This produced a total of 992,220,763 short reads. Whole-genome sequencing data are available in the Sequence Read Archive under BioProject PRJNA713582.

Quality of sequencing reads was evaluated using *FastQC* v0.11.8 (https://www.bioinformatics.babraham.ac.uk/projects/fastqc/). Reads were then mapped to our *de novo* reference assembly with *BWA-MEM* (Li et al., 2013) using default settings. Mapping statistics were calculated using *SAMtools* (Li et al., 2009) and custom scripts. Due to the low coverage of our dataset we conducted all our analysis in a genotype likelihood framework using the software *ANGSD* v0.930 (Korneliussen et al., 2014). When calculating genotype likelihoods in our analyses, we excluded: 1) triallelic positions (-skipTriallelic 1); 2) positions with base quality below 30 (-minQ 30); 3) reads with mapping quality below 30 (-minMapQ 30); 4) secondary and duplicate reads (-remove_bads 1); 5) reads with multiple best hits (-uniqueOnly 1); and 6) reads with one or both of the mates not mapping correctly (-only_proper_pairs 1).

### 2.6. Population genomics

We started our population genomic analyses by conducting a principal component analysis (PCA) on genotype likelihoods using *PCAngsd* with default parameters (Meisner & Albrechtsen, 2018). This program generated a covariance matrix between all individuals, which was used to estimate principal components and individual loadings with the function *prcomp* from *R* v3.6.3 (R Core Team, 2020). We complemented this approach by calculating individual admixture proportions with *NGSadmix* (Skotte et al., 2013). We ran the analysis for several values of *K* (2 to 6), imposing a minimum minor allele frequency of 0.05. We also calculated, for each pair of individuals, the relatedness index *r_xy_* (Hendrick & Lacy, 2015) from genotype likelihoods using *ngsRelateV2* (Hanghøj et al., 2019). PCA, admixture and relatedness analyses were run with the full dataset, imposing for each position (in addition to the filters outlined above) a minimum and maximum depths across all samples of, respectively, 10 and 200 (-setMinDepth 10, -setMaxDepth 200) and removing positions that did not have data in at least 15 individuals (-minInd 15).

Next, we investigated levels and patterns of genetic variation in migrant and resident white storks. For these analyses, we considered all 10 migrant storks that were not outliers in the PCA, and from the 40 non-outlier residents (see “Results”) we randomly sampled 10 individuals to avoid introducing biases related to sample size. Using *ANGSD*, we started by generating a maximum likelihood estimate of the unfolded site frequency spectrum and used this to calculate pairwise nucleotide diversity (π, (Nei, 1987)) and Tajima’s *D* (Tajima, 1989) in 200 kb non-overlapping windows. Unbiased nucleotide diversity was obtained by dividing the value obtained for each window by the total number of sites passing filters.

### 2.7. Selective sweep mapping

Population censuses detect a consistent and ongoing shift to residency in white storks from Portugal (Figure 1). If this phenotypic shift is mediated by selection acting on genetic variation, the genomes of resident white storks should exhibit significant deviations from neutrality. To detect signatures of selection in the genomes of residents (n = 43), we calculated several statistics that look at distinct but complementary properties of sequence data under selection. Specifically, we used: 1) Tajima’s *D* (Tajima, 1989), which detects deviations from neutrality by comparing the number of segregating sites and nucleotide diversity; 2) Fay and Wu’s *H* (Fay & Wu, 2000), which identifies recent selective sweeps by analysing the frequency distribution of derived alleles; and 3) the composite likelihood ratio statistic (CLR) from *SweepFinder2* (DeGiorgio et al., 2016), which uses the site frequency spectrum to identify loci affected by recent positive selection).

*ANGSD* was used to calculate *D* and *H* through the *thetaStat* module. Since *H* requires the identification of ancestral and derived alleles, we polarized polymorphic sites using sequence information from the closely related maguari stork (*Ciconia maguari*) obtained from NCBI’s SRA database (SRR9946656; Feng et al., 2020). We mapped these Illumina data to our reference genome (using the same approach as our own re-sequencing dataset; mapping rate 98.77%, coverage 21.97X), used *ANGSD* to generate a corrected haploid reference sequence (-doFasta 3), and finally used this corrected reference as an ancestral sequence when calculating the site allele frequency likelihood. For *SweepFinder2*, since the input file requires allele counts, we first used *ANGSD*’s single-read sampling method (-doIBS) to randomly subsample one allele per locus from each of the 43 individuals; these counts were used to generate an input file, and *SweepFinder2* was run under default parameters.

Results of the three tests were combined by calculating the de-correlated composite of multiple signals (DCMS, Ma et al., 2015), which takes into account the correlational structure of the variables to weigh their relative contributions to the combined score. Prior to this analysis, *D* values were multiplied by −1 so that higher values correspond to higher evidence of a selective sweep. Values of -*D*, *H* and CLR were each first converted into fractional ranks to create uniformly distributed probabilities, to which an inverse-normal transformation was applied. Normalised scores were transformed into Z-scores, and from these *P*-values were calculated assuming a normal distribution of the data. Spearman’s rank correlation coefficient was calculated for each pair of variables. Finally, correlation coefficients and *P*-values were used to calculate the DCMS score for each window (the top 1% outlier windows were considered as the most likely candidates for selection).

Although these approaches are suited to detect loci under selection in the resident white stork population, it is likely that a portion of the putative candidates of selection are not associated with the loss of migration, but rather with other local processes. To further filter out our list of potential candidates, we used *ANGSD* to calculate the fixation index (*F*_ST_) between migrants (n = 11) and residents (n = 43). To test for significant overlap between outliers of this statistic with those of DCMS, we used *SuperExactTest* (https://network.shinyapps.io/superexacttest; Wang et al., 2015). All genome-wide statistics were calculated in non-overlapping 50 kb windows; scaffolds smaller than 1 Mb were discarded, resulting in 22,766 windows that were shared between these analyses.

## 3. RESULTS

### 3.1. Individual consistency in migratory behaviour

GPS-tracking of 80 adult (≥ 4 years old) and 133 juveniles tagged as fledglings revealed extensive variation in migratory behaviour of white storks (Figure 2A, B and C) but a gradual decrease in the probability of migration with age (GLMM: −12.8 ± 2.6, *n* = 252, *P* < 0.001; Figure 2D). Despite the considerable inter-individual variability in migratory patterns of adult storks (Figure 2C), individuals tracked from 2 up to 7 consecutive years (*n* = 48) exhibited remarkable consistency in their migratory strategy over multiple years, revealing high intra-individual repeatability in the choice of the wintering region (*r* = 0.996, SE = 0.001, *P* < 0.0001, *n* = 161 bird-year comparisons, Figure 3). Only on three occasions did adult storks change migratory decisions; one bird in 2017 and two birds in 2021 remained in Iberia, after having migrated to Morocco in the previous year (Figure 3A). Overall, amongst tracked adults, only 19% migrated to Africa (wintering in Morocco or Sub-Saharan Africa) while 79% of the individuals remained in Iberia (Portugal and Spain), thus corroborating the national survey results (Figure 1C).

**Figure 2.**
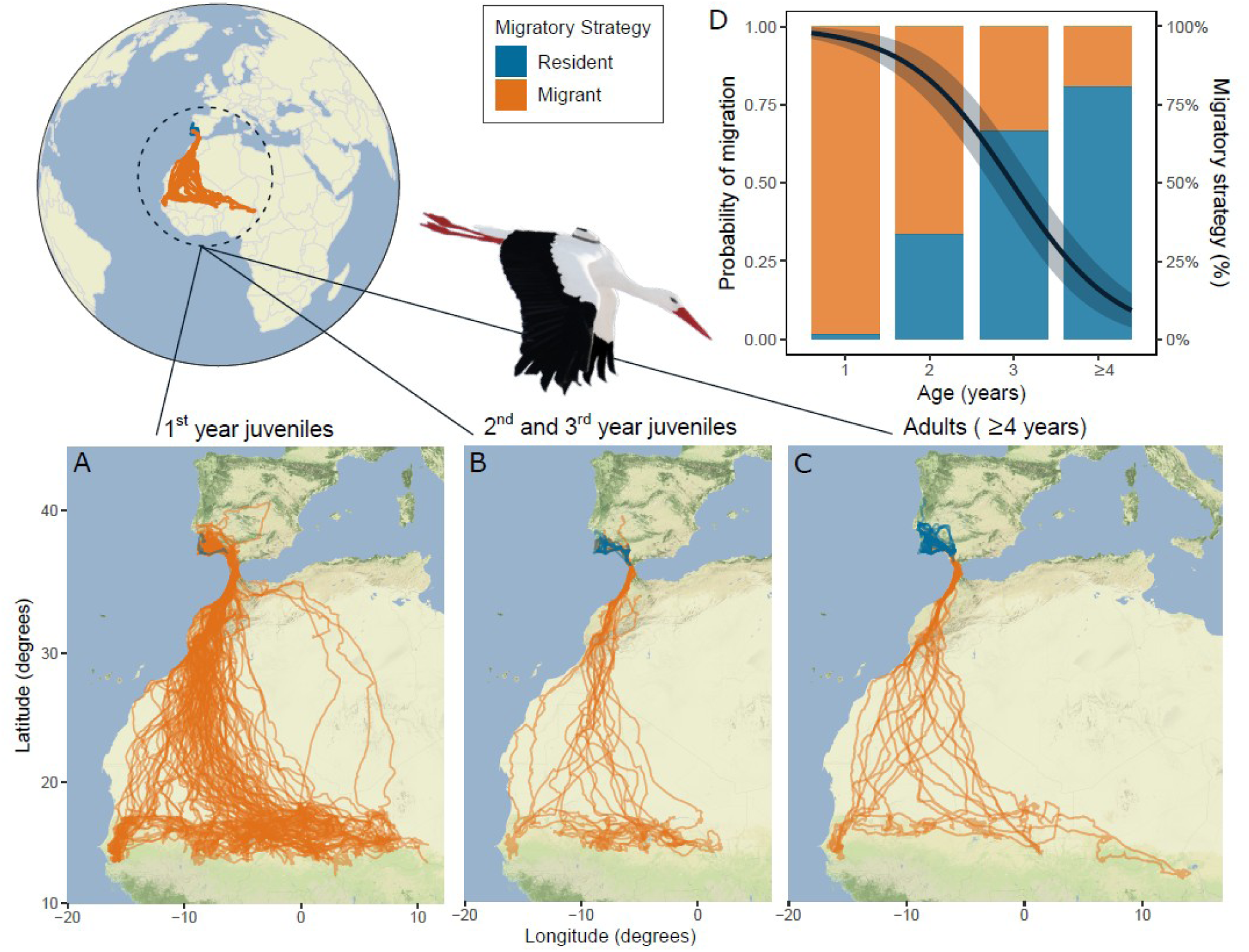
Migratory behaviour of GPS-tracked white storks from Portugal. Maps show GPS tracks of (A) First-year juveniles. (B) Second- and third-year juveniles. (C) Adults (≥ 4 years). Individual routes are coloured according to whether they were classified as migratory (orange, wintering in Africa) or resident, i.e., non-migratory (blue, wintering in Iberia). (D) Probability of migration by age. The regression line (black) was fitted using a binomial generalised linear mixed model testing the effects of age on the probability of migration and 95% confidence intervals (grey area) are shown. Columns reflect the percentage of migratory and resident individuals in each age class (*n* = 133, 24, 12, and 84 individuals with 1, 2, 3 and ≥ 4 years, respectively). 98% of all first-year juveniles migrated to Africa, but in their second and third years, the percentage of migratory individuals decreased to 67% and 33%, respectively. Amongst adults, only 19% of all tracked individuals crossed the Strait of Gibraltar. Three adult individuals that changed migratory strategy between consecutive years were not included.

**Figure 3.**
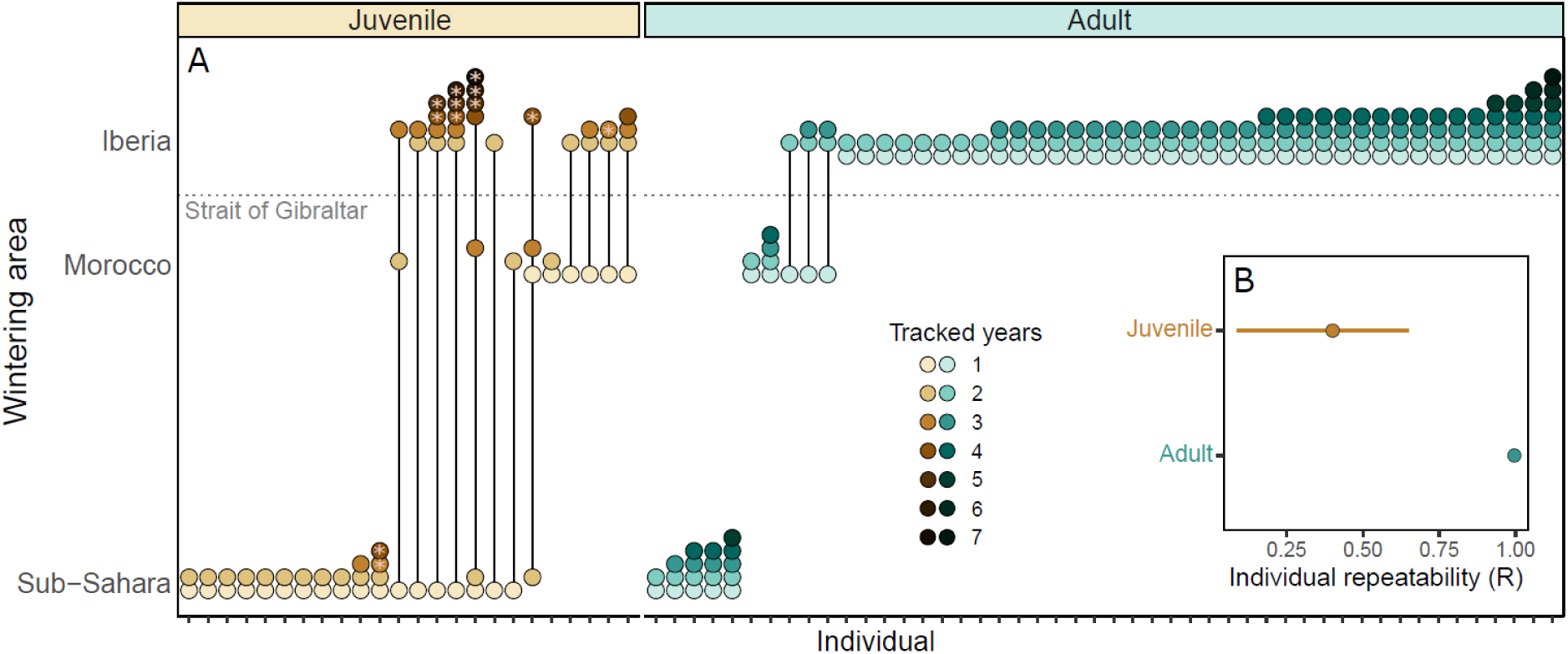
Consistency of white stork migratory behaviour across years. (A) Wintering areas of storks tagged as first-year juveniles (brown, *n* = 24) and adults (blue, *n* = 48) and tracked for multiple years. X-axis represent different individuals and dots represent annual wintering areas, coloured by the number of tracked years (lighter dots show younger birds). Lines indicate shifts in migratory behaviour in consecutive years. Horizontal dashed grey line represents the Strait of Gibraltar: dots above and below the line show resident (wintering in Iberia) and migratory (wintering in Africa) individuals. Asterisks represent the year in which the juvenile became a breeding adult. (B) Individual repeatability (R) in the choice of wintering grounds (minimum latitude reached in October) by first-year juvenile and adult (> 4 years) white storks tracked in multiple years. Whilst adults exhibited high intra-individual repeatability (*r* = 0.996, SE = 0.001, *P* < 0.0001), juveniles (excluding the breeding years) showed low repeatability (*r* = 0.401, SE = 0.14, *P* = 0.005) in the choice of the wintering region.

Contrarily to adults, juvenile white storks tracked until reaching sexual maturity (i.e., tracked for >1 and up to 3-4 years, *n* = 24) were not consistent in the choice of their wintering grounds in consecutive years (*r* = 0.401, SE = 0.14, *P* = 0.005, *n* = 59 bird-year comparisons, Figure 3). Amongst first-year juveniles, 98% crossed the Strait of Gibraltar towards their African wintering grounds, travelling shorter (13%) or longer (87%) distances to winter in Morocco or in Sub-Saharan Africa (Figs. 2B and 3A). Juvenile storks either decreased or maintained migratory distance (wintering at northern or similar latitudes) as they aged, with only one exception, a bird that wintered in Morocco and in the following year travelled to the Sahel, Figure 3A. Amongst young storks, the proportion of migrants decreased to 67% and 33%, in their second and third year of life, respectively. Moreover, in their third year, only 17% carried out long-distance movements to reach Sub-Saharan Africa (Figure 3A). Finally, all first-year juveniles tracked until the age of first breeding became consistent in migratory strategy as adults, wintering in Iberia or in Africa (*n* = 5 and *n* = 1 respectively, Figure 3A). These results show that the change in migratory behaviour in this long-lived species is associated to the immature stage (2 to 3-4 years), with the ultimate migratory strategy likely being acquired in the transition to adulthood, and thereupon maintained as storks establish themselves as breeders.

### 3.2. Population genomics of migrant and resident white storks

To assess the role of genetic variation in white stork migration, we started by assembling a *de novo* reference genome of the white stork using linked-read sequencing (Weisenfeld et al., 2017). This resulted in a 1.26 Gb reference sequence, comprised of 6,992 scaffolds (Table S1). A *BUSCO* search for highly conserved single-copy orthologues using an avian database revealed our genome to be highly complete: out of a total of 8.338 genes tested, 8.091 (97.1%) were present in the assembly. A lift-over of protein-coding gene sequences from the ruff genome annotation identified the coordinated for 17,535 genes (95.6% of the total 18,342 annotated ruff genes).

Following *de novo* assembly, we re-sequenced the genomes of 54 adult birds at 2.1X coverage (migrants and residents, Table S2), and investigated patterns of genomic variation in a sample of migrant and resident white storks. A principal component analysis (PCA, Figure 4A) indicated the presence of two clusters of samples along the first principal component but, most importantly, showed no genetic structure between samples according to their migratory behaviour in either PC1 (Mann-Whitney U test, *P* = 0.968) or PC2 (Mann-Whitney U test, *P* = 0.211). An admixture analysis indicates the same pattern, with migrant and resident individuals sharing ancestry to the same clusters along several values of *K* (Figure S3). The low overall genetic differentiation occurred despite no evidence of substantial relatedness between individuals in our dataset. Out of a total of 1,431 pairwise comparisons between samples, only 10 had an *r_xy_* index above 0.03125 (1/32), corresponding to a relatedness higher then second cousins (average ± standard deviation; *r_xy_* = 0.002±0.013). Concordantly with observations that migrant and resident individuals share the same ancestry and belong to a single interbreeding population, the average genome-wide nucleotide diversity (π) and a measure of the allele frequency spectrum of mutations (Tajima’s *D*) are similar between groups (average ± standard deviation; π_migrants_ = 0.085±0.045; π_residents_ = 0.083±0.041, *D*_migrants_ = 1.219±0.276; *D*_residents_ = 1.279±0.262, Figure 4B). Overall, our results indicate that the recent abrupt shift towards loss of migratory behaviour in white storks, which leads to large inter-individual differences in movement patterns, is not associated with differentiated populations within its Iberian breeding grounds.

**Figure 4.**
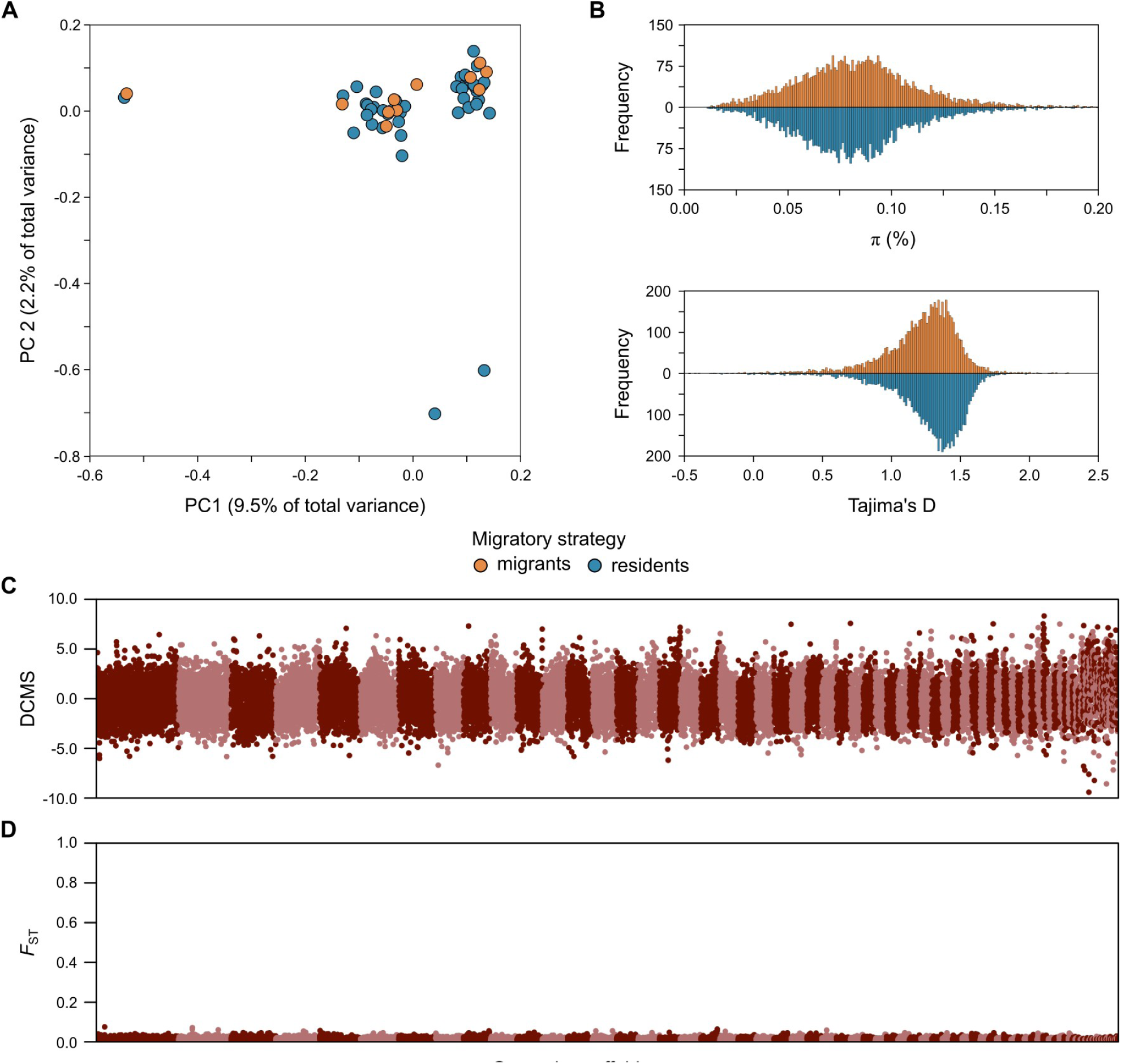
Population genomics of migrant and resident adult white storks. (A) Principal component analysis of genotype likelihoods. The percentage of variance explained by each component is given in parentheses. Individuals are coloured according to their migratory strategy. (B) Frequency distribution of genome-wide values of nucleotide diversity (π) and Tajima’s *D* of migrant and resident white storks, calculated in non-overlapping 200 kb windows. For π, a small number of windows above 0.2 were omitted to improve visualization. (C) Signatures of selection on the genome of resident white storks, measured by the de-correlated composite of multiple signals (DCMS), which summarizes patterns of Tajima’s *D*, Fay and Wu’s *H* and *SweepFinder2*’s composite likelihood ratio. (D) Genetic differentiation between migrant and resident white storks, measured by the fixation index (*F*ST). In panels (C) and (D) the statistics were calculated in non-overlapping 50 kb windows across the genome; colours indicate alternating genomic scaffolds. Despite differences in migratory strategy between individuals, the genomes of migrant and resident white storks were not differentiated and have similar levels and patterns of nucleotide variation.

To examine evidence of recent selection associated with loss of migratory behaviour, we scanned the genome of resident storks for signatures of selective sweeps using statistics that highlight different properties of genetic variation (Tajima’s *D*, Fay and Wu’s *H* and *SweepFinder2*’s CLR, Figure S4). As expected, overall correlation coefficients between the three tests had a wide range of variability (ρ*D-H* = 0.400; ρ*D-*CLR = 0.199; ρ*H-*CLR = −0.109). These should however converge in genomic regions under selection, so we summarized this information with the composite statistic DCMS (Figure 4C). The top 1% of the empirical distribution of DCMS (228 non-overlapping 50 kb windows) overlapped the full open-reading frame of 223 protein coding genes. None of these windows was an outlier in all the three statistics independently, and the majority represented isolated windows (197 independent regions, once adjacent windows were merged). The top outlier region (scaffold50:1,200,000-1,450,000), contained 4 genes, *EPHB6* (ephrin type-B receptor 6) and three T-cell receptor β chain genes. Other large outlier regions, containing three or more adjacent windows were located on scaffold10 (6,550,000-6,700,000; 2 genes), scaffold17 (23,550,000-23,750,000; 14 genes), scaffold19 (19,700,000-19,850,000; 3 genes) and scaffold49 (1,450,000-1,600,000; 1 gene).

While these genomic regions may represent evidence for selection acting on the resident stork population, it is possible that they reflect selection on traits not associated with migration. To test this, we explicitly calculated genome-wide differentiation (*F*_ST_) between migrant and resident storks. Strikingly, we failed to find substantial differences between individuals of the two groups at any locus across the genome (Figure 4D). The highest *F_ST_* values were also scattered throughout the genome (182 independent regions, once adjacent windows were merged) and, more importantly, were of very low magnitude. To illustrate this, the top window had an *F*_ST_ of 0.076, and only 15 windows (0.07% of the total number) had an *F*_ST_ above 0.05. The average *F*_ST_ in the top 1% DCMS windows (*F*_ST_ = 0.015) was also similar to the genome-wide average (*F*_ST_ = 0.014). A comparison of the top DCMS outliers with the top *F*_ST_ outliers indicated a general lack of concordance (Figure S5), since only four genomic windows were simultaneously 1% outliers in both statistics, an overlap that wasn’t higher than expected by chance (*P* = 0.196). Of these four windows, three were placed on scaffold3 (no gene was located 100 kb either side of these windows) and a window on scaffold40 (overlapping genes *HAO2* and *3BHSD*); patterns of *F_ST_* between the two groups in these genomic regions do not conclusively suggest an association (Figure S6). This likely lack of a genetic association in our study does not preclude a scenario of selection acting on an unassembled region of the genome (unlikely given our genome is highly complete), or that the shift could be associated with widespread polygenic selection characterised by very small changes in allele frequency (which would require a very large sample size to detect). However, taken together with our tracking data, these results support a scenario in which developmental effects are the major mechanisms explaining the rapid loss of migration in white storks, albeit with potential minor contributions from phenotypic flexibility or selection on genetic variation.

## 4. DISCUSSION

Unveiling if, and how, natural populations respond to ongoing human-driven environmental changes is a topic of central importance in evolutionary ecology and conservation. Given the potential for animal migration to alter ecological networks worldwide, understanding the roles that selection and plasticity play in shaping migratory strategies will be key to obtaining a full comprehension of how this life history trait can change in response to environmental changes. Our results show that rapid and drastic population-level changes in avian migratory behaviour, including the loss of migration itself, can arise through generational shifts, with young recruits as the agents of such changes. In white storks, the documented shortening of migratory distances, and even the loss of migratory behaviour (Archaux et al., 2004; Catry et al., 2017; Cheng et al., 2019; Flack et al., 2016; Rotics et al., 2017), is likely occurring through inter-generational processes driven by the recruitment of non-migratory young storks into the population. This could represent an effective mechanism for rapid acclimation to anthropogenic environmental change.

Individual long-term tracking of adult white storks revealed high within-individual consistency in migratory strategy (either migratory or non-migratory), thus not supporting recent findings reporting phenotypic flexibility as the mechanism driving population-level changes in avian migratory behaviour (Conklin et al., 2021; Fraser et al., 2019). Between 2016 and 2022, only three adult storks shifted migratory strategy (3 out of 113 potential transitions), from short-distance migration to Morocco to residency in Iberia. This observed adult flexibility (0.4% per year) would not be enough to explain the magnitude of observed changes in migratory behaviour of the Portuguese stork population in the last 25 years, where 50-64% of the population became non-migratory (Figure 1C). One can argue that the high consistency in migratory strategy amongst resident birds does not derive from an inflexible phenotype, but instead from the favourable and stable environmental conditions in Iberia during the non-breeding season (due to high food availability in landfills). However, if adults are indeed flexible in their migratory behaviour, we would expect long-distance migrants, that are exposed to annual variability in wintering conditions in the Sahel, to alternate between migratory strategies. Instead, we found an exceptionally high within-individual consistency in migratory behaviour, even (and particularly) in long-distance migrants over multiple consecutive years (R = 1), suggesting that other mechanisms must be involved in the loss of migratory behaviour. Contrarily to adults, most tracked juveniles experienced a shift towards residency throughout their development, before reaching sexual maturity (3-4 years), suggesting that through inter-generational shifts in the frequency of non-migratory recruits in the population, a population can change from mostly migratory to mostly resident. This is supported by a population viability analysis that indicates that a comparable phenotypic shift at the population level can be attained by a conversion rate of just 10% of migratory juveniles to resident adults (Supplementary text, Figure S7, Table S3, Table S4).

As social migrants, with naïve birds requiring guidance from older individuals to successfully reach their wintering areas, we would expect that most first-year storks would remain in Iberia, alongside non-migratory adults. Yet, the species innate migratory program likely drives all first-year juveniles to complete an initial migration to Africa (Chernetsov et al., 2004; Mayr, 1952). On their second- and third-years, environmental, physiological, individual experience, or social cues could decrease the propensity to migrate and override the initially expressed migration program, leading to irreversible adult phenotypes (Åkesson & Helm, 2020; Gill et al., 2019; Méndez et al., 2021). Indeed, whilst 98% of first-year juveniles migrated to Africa, the proportion of migratory individuals decreased to 67% and 33% in their second and third years. Moreover, first-year juveniles tracked beyond age of first breeding maintained their migratory strategy through adulthood (five juveniles became residents and one remained migrant, wintering in the Sahel), adding to the findings that migratory strategy is not fully behaviourally flexible, becoming locked at a certain stage of individual development.

Although for many species the propensity to migrate long distances has a strong genetic component (Liedvogel, 2019) that could be under selection due to environmental change, we did not find conclusive support for the hypothesis that selection drives loss of migration in the white stork. It should, however, be noted that our limited sample size may preclude a definitive answer on the role of selection if it is associated with soft sweeps on multiple loci of small effect (Hermisson & Pennings, 2017). In any case, the lack of a clear signal across several independent statistics is indicative that a recent sweep based on a large effect locus, as found in previous genomic studies, is unlikely. While the innate migratory program presents an opportunity for selection to drive changes in migratory strategy, plasticity should provide a faster mechanism for adaptation. An association between changes in migratory strategy and large-effect genetic variation is thus more likely to be found in smaller birds (nocturnal solitary migrants) (Delmore et al., 2016, 2020; Lundberg et al., 2017; Sanchez-Donoso et al., 2022; Toews et al., 2019), which are typically short-lived and thus less likely to modulate migration based on ontogenetic effects or experience (Pulido, 2007).

The observed shortening of migration distance and suppression of migratory behaviour in white storks has been associated with an increased year-round food availability from landfills (Catry et al., 2017; Cheng et al., 2019; Flack et al., 2016). Recent studies showed that foraging on landfill waste is a time- and energy-saving strategy that enables storks to reduce their movement and foraging efforts (Soriano-Redondo et al., 2021), thus facilitating their survival through the winter season. Indeed, overwintering in North Africa (Morocco) and Europe, where landfills and rubbish dumps provide high abundance of food, has been shown to enhance white storks’ juvenile survival (Cheng et al., 2019; Rotics et al., 2017), likely contributing towards the manifestation of non-migratory behaviour. For immatures, the decision to stay in Iberia and no longer migrate to their sub-Saharan wintering grounds could also be strengthened by social learning (Byholm et al., 2022; Mueller & O’Hara, 2013; Teitelbaum et al., 2016), in which young storks acquire information from the increasing number of experienced conspecifics overwintering on these food waste disposal sites. Despite the evident benefits, foraging on landfill waste is also likely to exacerbate intraspecific competition (Martins et al. 2024, Soriano-Redondo et al., 2021). Thus, social interactions, mediated by each individual’s foraging proficiency and experience, could function to promote or suppress migration of dominant and outcompeted individuals, respectively (Campioni et al., 2020; Grecian et al., 2018). Finally, migratory propensity could also be altered during maturation through carry-over effects of differential migratory performance. Increased flight costs have been shown to be an important proximate cause of juvenile mortality during migration (Rotics et al., 2017) and poor individual performance during first migrations could suppress migratory behaviour in following years. Regardless of the driver behind shifts in migratory behaviour during juvenile development, disproportionate changes in survival rates of migratory and non-migratory recruits could, through generational shifts, accelerate the white stork population turnover towards residency. In the short term, if environmental conditions continue to favour non-migratory individuals, the white stork population is likely to change towards full residency. Yet, future waste reduction initiatives planned by the European Union (Soriano-Redondo et al., 2021) might revert this tendency in the medium to the long-term and generational shifts could again allow white storks to track future changes.

Our study expands on earlier findings on the evolution and development of bird migratory behaviour in three key aspects. First, we offer additional evidence that these developmental processes can lead not only to phenological and range changes in migratory movements (Gill et al., 2019; Verhoven et al; 2018, 2021), but also the undertaking of long-distance migration itself. Second, we present evidence that genetic variation, in particular in large-effect loci, is not associated with this shift. Third, we employed a large-scale longitudinal study throughout the earlier years of white storks, pinpointing the developmental interval during which dramatic changes in behaviour take place. By allowing organisms to develop phenotypes adjusted to the conditions that adults will experience, developmental plasticity can provide, through generational shifts, a fast mechanism for long-lived species to adapt to novel ecological opportunities within the lifespan of individuals and the topic is receiving increased attention (Åkesson &. Helm, 2020; Gill et al., 2019). Future research should thus focus on identifying the specific developmental mechanisms that drive migratory traits during ontogeny (e.g., flight efficiency, migratory performance, access to food resources at landfills), to further increase our understanding about species adaptation to environmental change and the associated implications for their conservation.

## Data accessibility

Chromium sequencing data for reference genome assembly and whole-genome re-sequencing data are available in the Sequence Read Archive (www.ncbi.nlm.nih.gov/sra) under BioProject PRJNA713582. The reference assembly sequence is deposited in GenBank (www.ncbi.nlm.nih.gov/genbank), under accession GCA_030584885.1.

## Authors’ contribution

I.C., A.M.A.F., F.M. and M.A. coordinated and performed collection of ecological and tracking data from white storks. M.C., S.A. and C.I.M. coordinated and performed collection of genetic data. P.A., M.C. and C.I.M performed genetic data analyses. I.C., M.A. and A.M.A.F. performed ecological and tracking data analyses. P.A. and I.C. wrote an initial version of the manuscript, which had input from all other authors.

## Acknowledgements

The authors thank Carlos Pacheco, Bruno Herlander and Katharine Rogerson for the help with white stork captures, Andrea Soriano-Redondo for contributions to management of tracking database, João Paulo Silva for help with programming the tracking devices and all volunteers that participate in the white stork census. Rui Faria, Paulo Pereira, Stephen Sabatino and Ricardo Jorge Lopes provided useful suggestions on an earlier version of the manuscript.

## SUPPLEMENTARY TEXT

### 1. Non-breeding surveys and estimates of resident white storks in Portugal

The number of resident white storks was assessed from the number of individuals counted during the non-breeding surveys, performed from mid-September to early October. This period was chosen because most migratory individuals cross the Strait of Gibraltar towards their African wintering grounds between July and early September (Fernández-Cruz et al., 2005; Soriano-Redondo et al., 2020); indeed, among our tracked adult and juvenile storks, autumn migration started between 7^th^ of July and 4^th^ September (median date = 5^th^ August, SD = 17 days, *n* = 75; Acácio et al. 2022). The pre-nuptial return migration to the breeding areas starts in November (Fundación MIGRES, pers. comm.) and none of our GPS-tracked storks arrived in Portugal before November. However, counts performed during this period could overestimate the number of resident storks due to the inclusion of storks of non-Portuguese origin that overwinter in Portugal. Nonetheless, most of the storks from the Central and Northern European populations migrating through or spending the winter in Iberia, seem to select further eastern routes/wintering areas, and the ones traveling to Portugal, seem to arrive mostly from mid-October onwards. Indeed, publicly available tracking data on Movebank studies with available visualization of tracks, show that only 9 out of 342 white storks migrating through/to Iberia visited Portugal (number of storks tagged in Austria = 5, France = 29, Germany = 236 and Spain = 72). Careful observation of individual tracks and data available on the Movebank Data Repository shows that only 1 out of the 9 storks visiting Portugal was present during the survey period (15 Sep-15 Oct), thus representing less than 1% (0.003%) of all tracked individuals.

Resights of ringed white storks at Portuguese landfill sites during monthly counts in 2019 and 2020 also support the hypothesis that most foreign individuals (from other European countries) arrive from late October (Figure S2). Overall, the inclusion of non-Portuguese storks in the censuses of resident storks should be negligible and likely compensated by the absence of Portuguese resident storks wintering in southern Spain (observed from GPS tracked individuals).

*Movebank studies:* Cicognes de Loire-Atlantic; Ciconia ciconia Sudewiesen; Cicognes de Saintonge; HUJ MPIAB White Stork E-obs; HUJ MPIAB White Stork GSM 2013; HUJ MPIAB White Stork GSM E-obs; Life Track White Stork Bavaria; Life Track White Stork Catalonia; Life Track White Stork Loburg 2022; Life Track White Stork Oberschwaben; Life Track White Stork Rheinland-Pfalz; Life Track White Stork Sarralbe; Life Track White Stork Spain Donana; Life Track White Stork SW Germany; Life Track White Stork SW Germany Care Centre Release; Life Track White Stork SW Germany CASCB; Life Track White Stork Vorarlberg; MPIAB Argos white stork tracking (1991-2022); White Stork Affenberg releases MPIAB; White Stork Loburg 2014.

*Movebank data repository accessions:* doi:10.5441/001/1.v1cs4nn0; doi:10.5441/001/1.c42j3js7; doi:10.5441/001/1.4192t2j4; doi:10.5441/001/1.ck04mn78; doi:10.5441/001/1.71r7pp6q

### 2. Population viability analysis

White stork numbers increased in Portugal in the last few decades (Catry et al., 2017) following a period of population declines observed in the first half of the 20th century (BirdLife International, 2016). In Portugal, census data show the breeding population increased 350%, from 3302 breeding pairs in 1994 to 11691 breeding pairs in 2014. During this period, the proportion of resident storks increased from 18% to 61.7%. A subsequent winter census in 2020 counted 19,282 white storks wintering in Portugal, corresponding to 67.6-82.5% of the breeding population.

We used a population viability analysis (PVA), incorporating biological and environmental variables, to explain the observed changes in the number of migrant and resident white storks in the last decades, and compared two scenarios: (1) populational shift towards residency explained solely by differences in demographic parameters between resident and migrant storks, with no individual changes in migratory behaviour; (2) populational shift towards residency explained by the loss of migratory behaviour during ontogeny, considering the conversion of juvenile migrants to residents, followed by consistency in migratory strategy in the adult life stages, as hypothesized in this study.

#### Methods

We predicted the trajectories of the migrant and resident populations from 1994 to 2020 using the average demographic parameters for our monitored white stork population (authors’ unpublished data, Soriano-Redondo et al., 2023). When parameters were not available for the Portuguese white storks, we used the average from Western European white stork populations summarised in Mayall et al. (2023). The demographic parameters used and the two scenarios are summarised in Table S3. The demographic trajectories for both migratory and resident populations were obtained using *Vortex* 10.5.20 (Lacy, 2019). *Vortex10* is an individual-based simulation model in which populations are subjected to a set of deterministic environmental, demographic, and genetic stochastic events (Brook et al., 1999). The rationale for the different parameters that were applied was as follows:

a. **Initial population size**: The initial population size was set to 5,416 migratory and 1,188 resident white storks, as obtained during the first breeding and wintering census in 1994 (Catry et al., 2017). We ran the demographic models for 26 years, from 1994 to 2020, as this was the year when the last census was performed and for which we have demographic data, reducing the influence of uncertainties in parameter estimations.
b. **Dispersal**: We modeled two scenarios. In scenario 1, we investigated if the differences in demographic parameters of the migratory and resident storks could drive the observed population changes (i.e., no individual changes in migratory strategy, hence no dispersal between the two populations). Scenario 2 included 10% of dispersal from the migratory to the resident population, during the storks’ 2nd and 3rd years of life, simulating the observed changes in the migratory behaviour of juvenile storks. The current observed rates of dispersal reported in this study are higher, yet these rates are likely to have changed through time, hence 10% was considered an average value over 26 years modelled.
c. **Reproductive system**: White storks are socially monogamous, with high nest fidelity (Barbraud et al., 1999) and moderate levels of extra-pair paternity (Turjeman et al., 2016). Therefore, we described the mating system as long-term monogamous relationships in *Vortex10*. The maximum age of breeding for both males and females was set to 30 (Mayall et al., 2023), with one brood per year. Age of first breeding can vary but generally starts in the 3rd year of life (Barbraud et al., 1999; Hancock et al., 1992; authors’ unpublished data); there are some recorded cases of two-year-old storks attempting to reproduce, although rarely successfully (Barbraud et al., 1999).
d. **Reproductive rate**: We assumed that 100% of adult females aged three years and older would attempt to breed with a 10% standard deviation due to environmental variation. Based on the average information obtained from 2016 to 2020, 95% of the females successfully fledged young at a rate of 1.7 (SD=0.45) fledglings per successful nest (authors’ unpublished data). The maximum progeny per brood was set to four (authors’ unpublished data). There were no differences in reproductive rates between the two populations as reported in Soriano-Redondo et al. (2023).
e. **Mortality rates**: Mortality rates vary significantly with age, both for migrants and resident storks. In the first year of life, most storks migrate, hence mortality is the same for both populations. First-year storks have high mortality rates due to low foraging experience and less efficient flight strategies (Kanyamibwa et al., 1990; Rotics et al., 2016). Collisions with powerlines can also contribute heavily to post-fledging mortality (Tobolka, 2014). Juveniles (ages 0–1) were assigned a mortality rate of 65.1% (SD due to environmental variation = 10) based on a mean value weighted by sample size for the western European stork population (extracted from Mayall et al. 2023). In the second year of life mortality declines and was set at 22.2% (SD due to environmental variation = 3) for migratory and 17.7% for resident storks based on average values obtained from Mayall et al. (2023). The mortality of migratory birds 3+ was set at 11% (SD due to environmental variation =3) for residents and 9% (SD due to environmental variation =3) for the migratory birds (Soriano-Redondo et al., 2023).
f. **Genetics**: The white stork’s long lifespan and slow generation time may result in the presence of deleterious genetic effects which may be hidden. *Vortex10* considers the lethal equivalent (LE) as a unit of deleterious genetic variation that when dispersed amongst a group of individuals could result in the mortality of specific individuals (Kalinowski & Hedrick, 1998). Due to high dispersal and strong population growth rates, all models were considered to have 0 LE. There is evidence to suggest that European white storks did not lose a significant amount of genetic diversity following their twentieth-century decline, suggesting that any negative repercussions from inbreeding depression are unlikely to have hindered the Portuguese populations (Shephard et al., 2013).
g. **Carrying capacity**: The carrying capacity was set at 35,000 which is well above the population sizes observed at present. This value was selected to not limit population growth rates. There were no other management measures included in the simulations.

#### Results

The demographic parameters used to determine the population trajectories simulated well the increase in white stork numbers observed over the 26 years (Figure S7), the overall white stork population increased to 28,842-31,862 individuals, according to scenarios 1 and 2 respectively (Table S4). Both simulated populations are similar to the current white stork population size (28,500 storks). However, in scenario 1 (no changes in migratory behaviour) the percentage of resident storks at the end of the 26 years, in 2020, (26.9%, Figure S7) is significantly lower than the observed percentage in our studied population (67.6 – 82.5%, Figure 1C in the manuscript), while in Scenario 2 (including 10% dispersal from the migratory to the resident population during ontogeny), the number of residents increased sharply and the percentage of resident individuals (69.8%, Table S4) is similar to the observed percentage in our studied population.

These simulations show that the ongoing changes in migratory behaviour in the Portuguese white stork population would not be possible without the conversion of migratory to resident storks. This conversion was only observed during ontogeny (2-3 years), as adults were consistent in their migratory strategy. Thus, we provide additional evidence that the observed turnover in the population migratory traits is occurring through generational shifts (with migratory juveniles settling as resident adults in the metapopulation).

## SUPPLEMENTARY FIGURES

**Figure S1.**
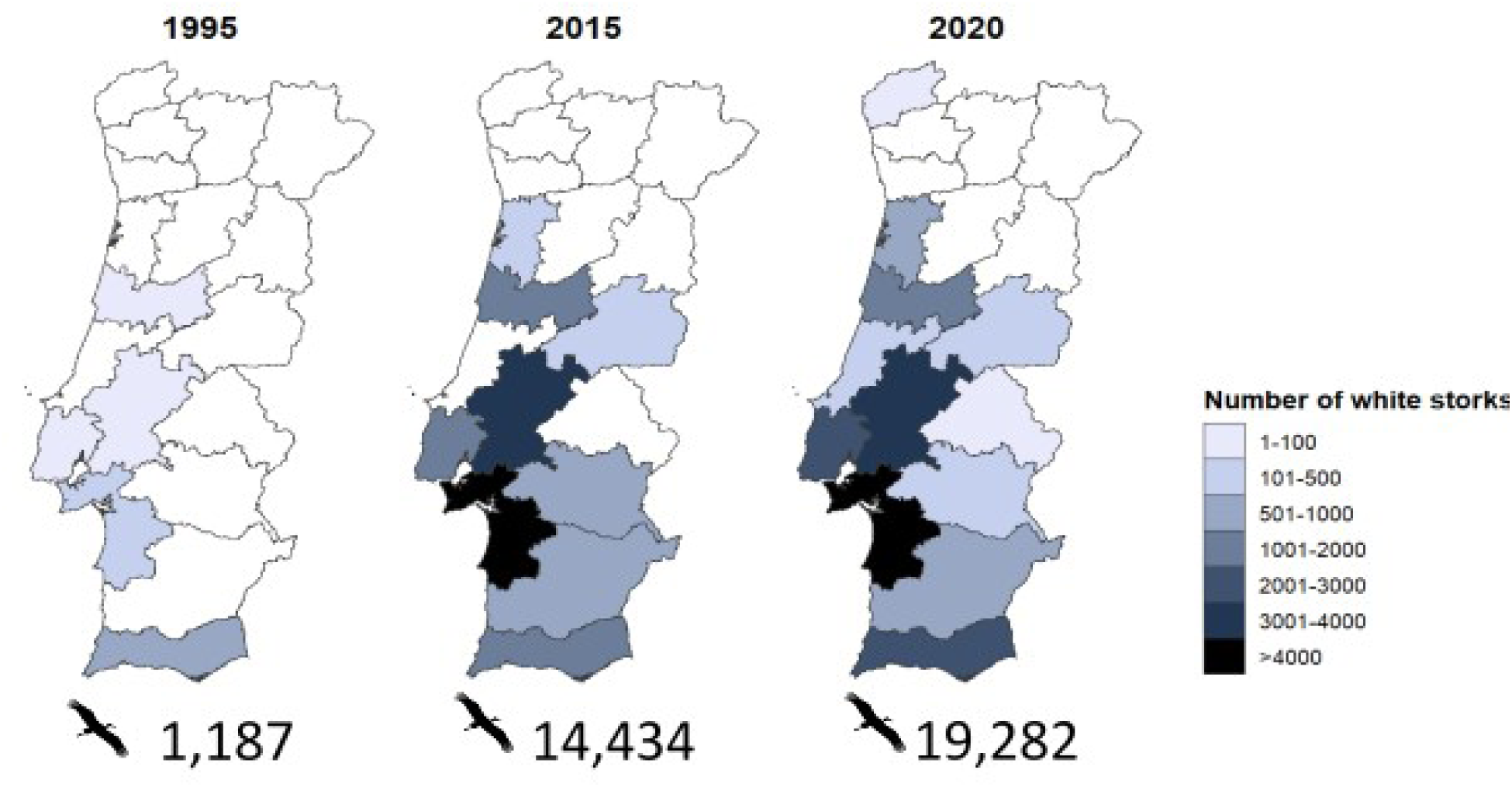
Number and distribution of wintering white storks in Portugal during the last 25 years. The total number of individuals counted during the non-breeding census in 1995, 2015 and 2020 is shown below the respective map.

**Figure S2.**
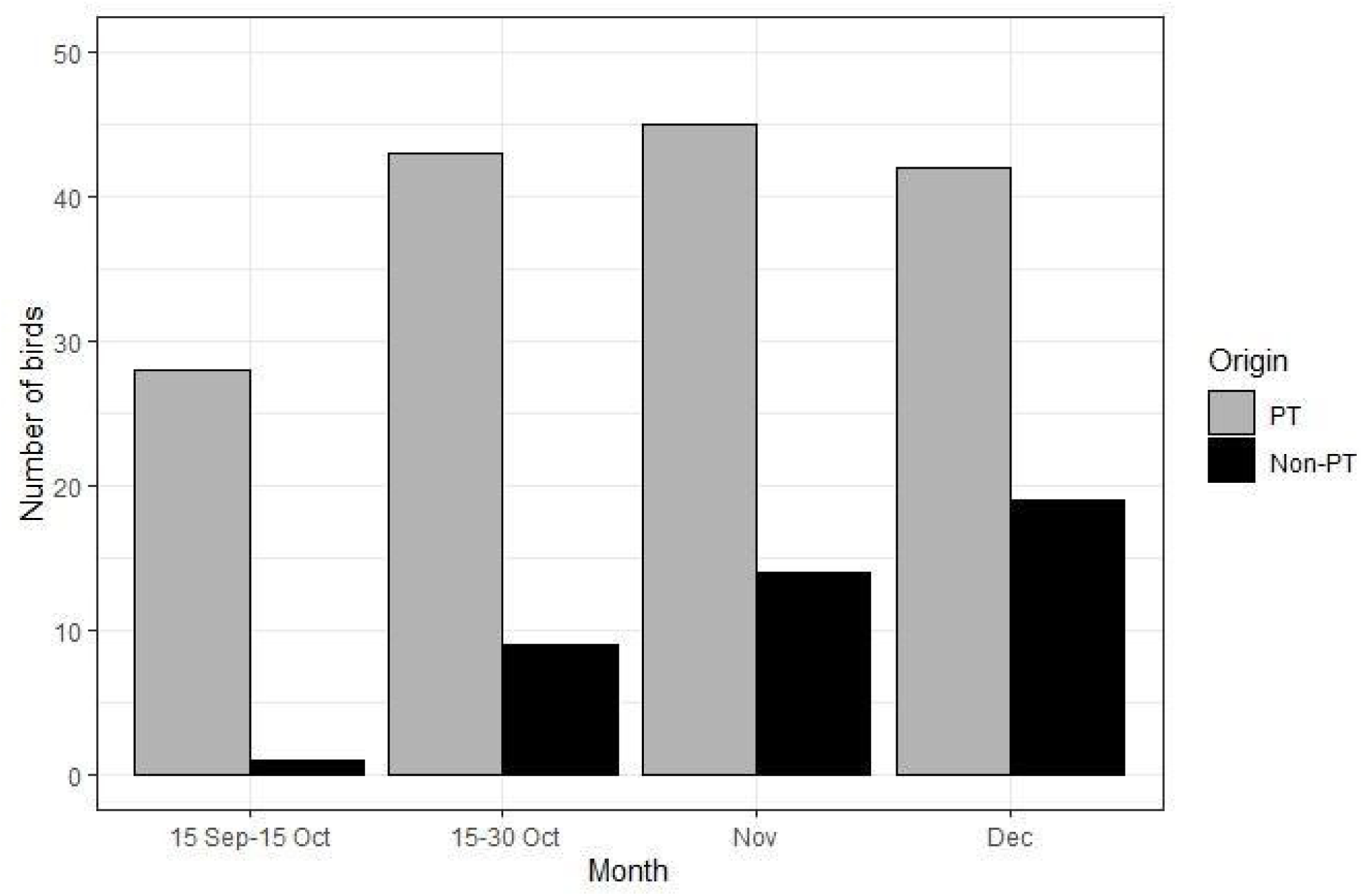
Number and origin (PT - Portuguese, Non-PT - other countries) of white storks resighted at four Portuguese landfill sites from September to December 2019-2020 (mean number of storks in landfills = 5100, 4850, 5075 and 3650 in September, October, November, and December, respectively). These data indicate that the overwhelming majority of birds detected during non-breeding surveys (carried out before the 15^th^ of October) were non-migrants of local origin.

**Figure S3.**
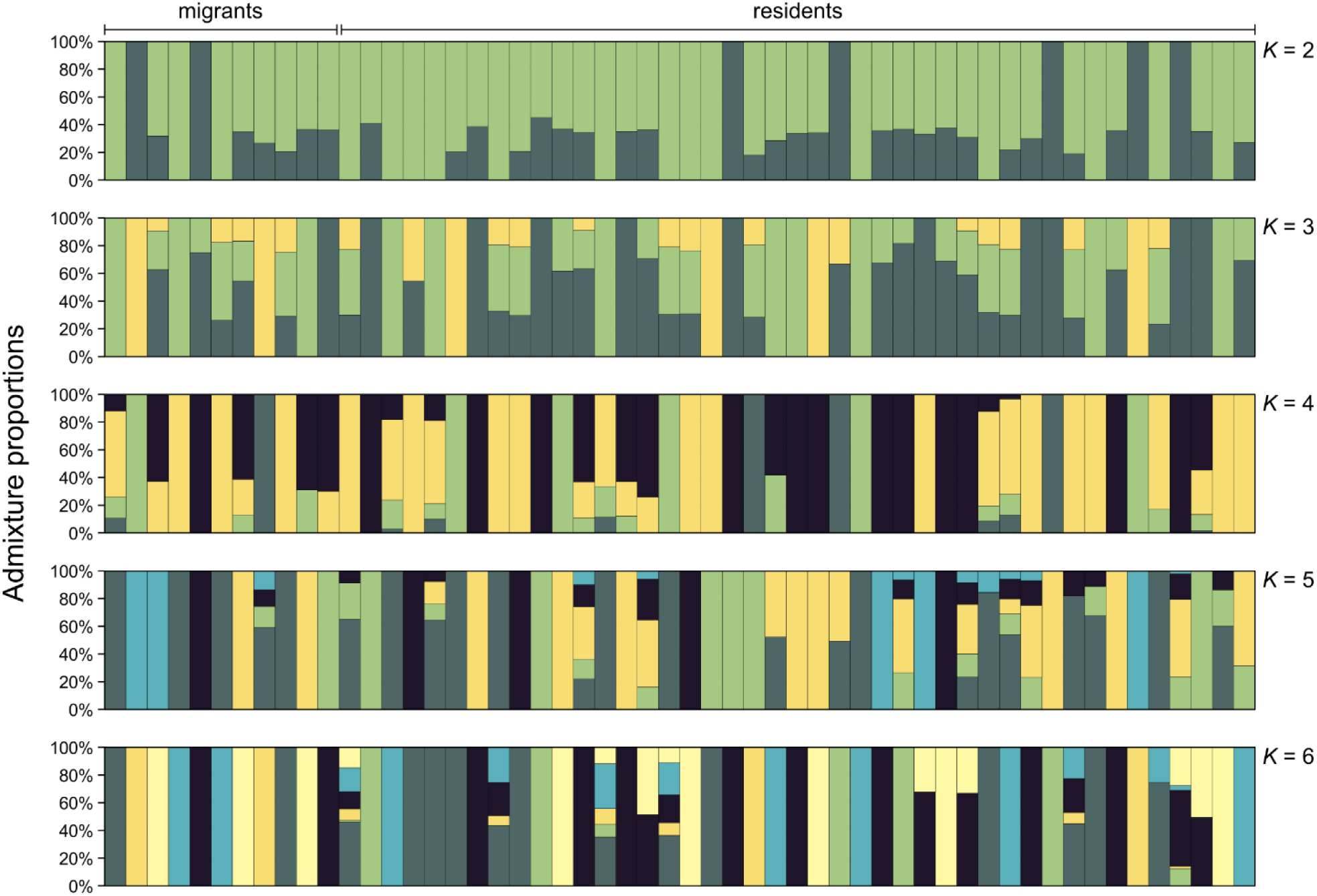
Individual admixture proportions for migratory (n = 11) and resident (n = 43) white storks, calculated using *NGSadmix*. Results for several values of K are shown (*K* = 2 to *K* = 6).

**Figure S4.**
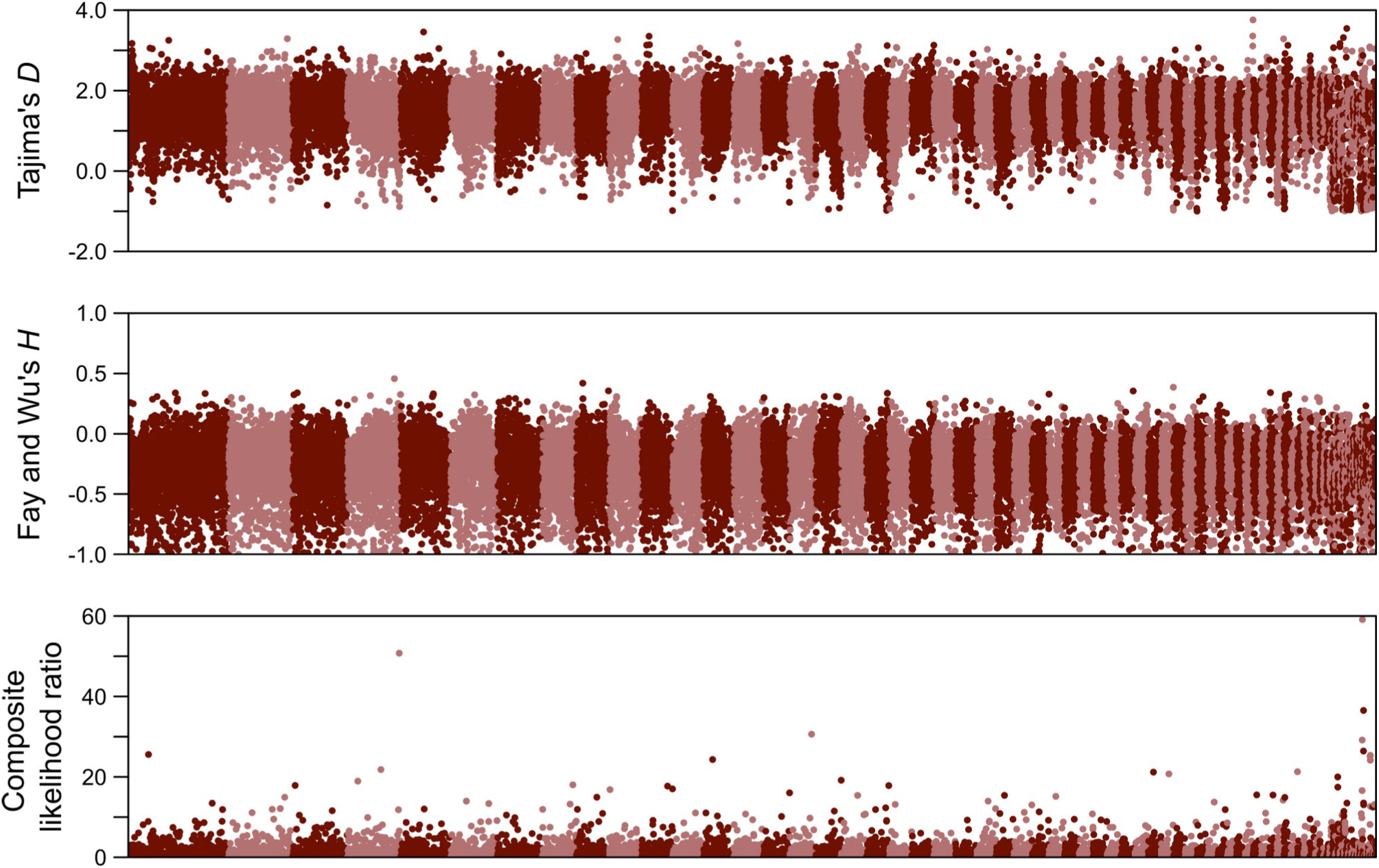
Manhattan plots with genome-wide scans for signatures of selection in resident white storks. Tajima’s *D* (top) tests for deviations from neutrality by comparing the number of segregating sites and nucleotide diversity; Fay and Wu’s *H* (middle) identifies recent selective sweeps by analysing the frequency distribution of derived alleles; the composite likelihood ratio statistic (CLR, bottom) from *SweepFinder2*, which uses the site frequency spectrum to identify loci affected by recent positive selection. The three statistics were calculated in non-overlapping 50 kb windows. Colors indicate alternating genomic scaffolds.

**Figure S5.**
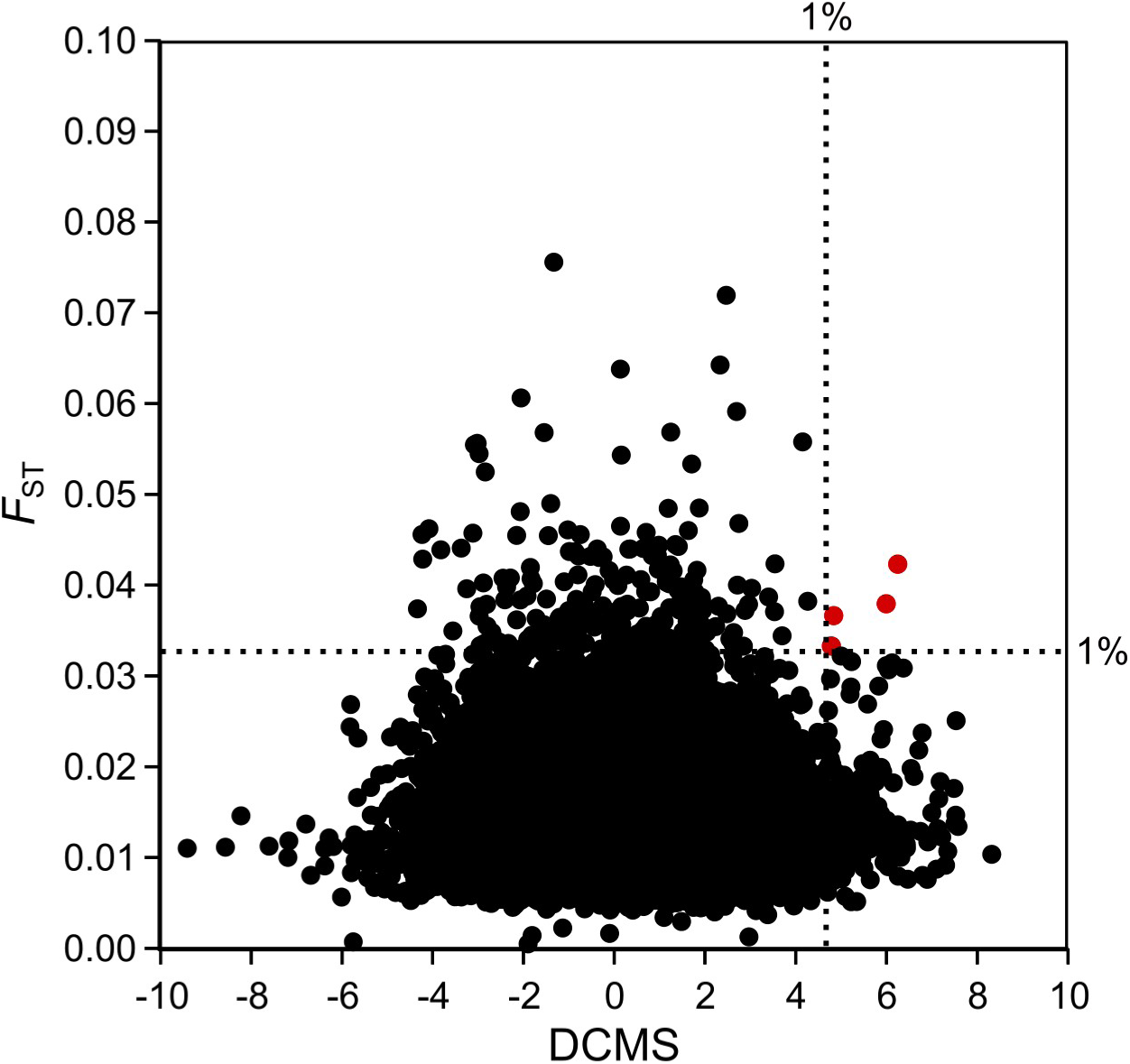
Comparison between values of genetic differentiation (fixation index, *F*ST) and the de-correlated composite of multiple signals (DCMS, summarizing patterns of Tajima’s *D*, Fay and Wu’s *H,* and *SweepFinder2*’s composite likelihood ratio), for 22,766 genomic windows of 50 kb (non-overlapping). The four windows that are within the top 1% of both statistics are coloured in red.

**Figure S6.**
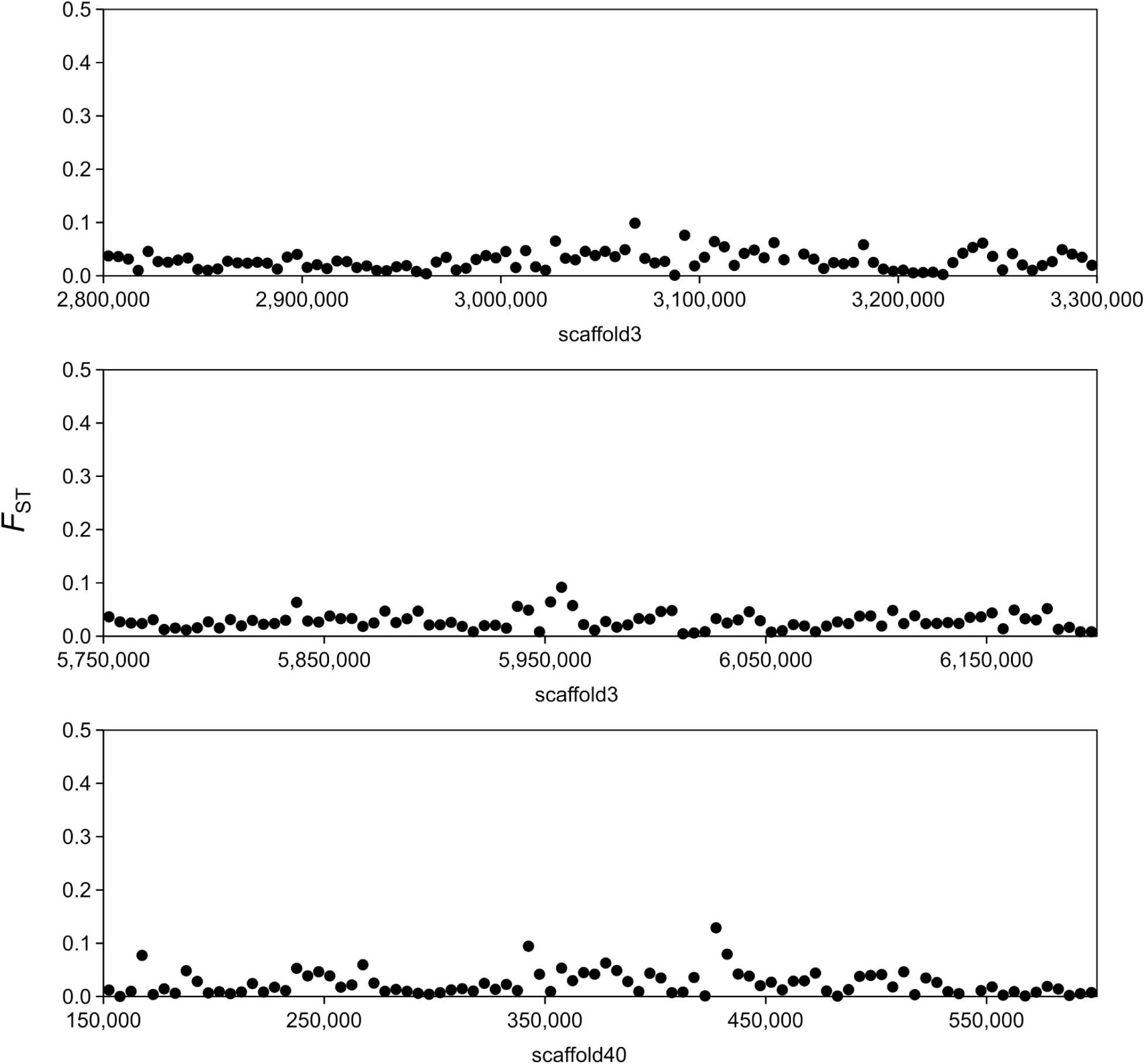
Genetic differentiation (fixation index, *F*ST) between migrant and non-migrant white storks, at the three genomic regions that were 1% outliers in the *F*ST and DCMS statistics (corresponding to the four windows in Figure S5). Each dot corresponds to a 5 kb non-overlapping window.

**Figure S7.**
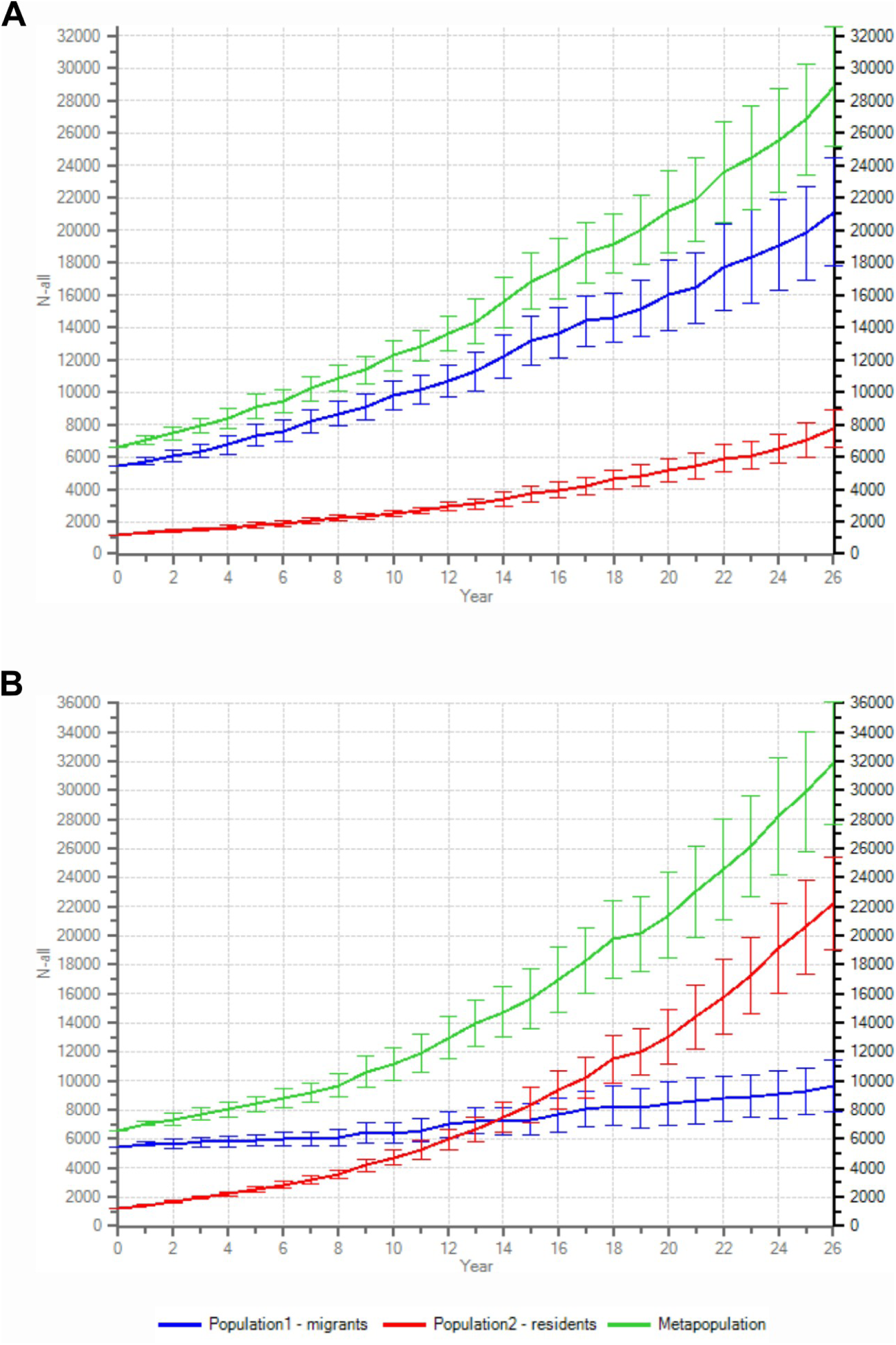
*Vortex10* simulations of white stork population trajectories obtained for 26 years from 1994 to 2020. The demographic parameters used are listed in Table S3. In scenario 1 (A) migratory and resident individuals do not shift migratory strategy. In scenario 2 (B) 10% of the migratory white storks (in the second and third year of life) shift to a resident strategy. The migratory population trajectory is shown in the blue line, the resident population is represented by the red line, and the metapopulation is represented in green, 95% confidence intervals are shown.

## SUPPLEMENTARY TABLES

**Table S1.** Summary statistics for the *de novo* genome assembly for the white stork *Ciconia ciconia* (Ccic_1.0). The assembly was done using 10X Genomics’ Chromium linked read technology. The sample used for the assembly was an adult female, resident in Iberia (ring number MR09149, CEMPA). The reference sequence is deposited in GenBank under accession GCA_030584885.1.

|  | **Ccic_1.0** |
| --- | --- |
| Assembly size (Gb) | 1.26 |
| Scaffold N50 (Mb) | 24.5 |
| Contig N50 (kb) | 300.8 |
| Total number of scaffolds | 6,992 |
| Number of scaffolds >10 kb | 890 |
| Number of scaffolds >100 kb | 165 |
| Largest scaffold (Mb) | 94.6 |
| GC content (%) | 42.1% |
| Complete BUSCOs reference (%) | 8,091 (97.1%) |
| Fragmented BUSCOs reference (%) | 49 (0.6%) |
| Missing BUSCOs reference (%) | 198 (2.3%) |

**Table S2.** Summary statistics of the whole-genome re-sequencing dataset.

| Sample | Ring number<br>(color ring) | Phenotype | Number<br>of reads | % properly<br>paired reads | % reads<br>MQ>=30 | Mean depth<br>of coverage |
| --- | --- | --- | --- | --- | --- | --- |
| whitestork_01 | MR09459<br>(70+) | resident | 2,832,800 | 95.5% | 92.0% | 0.3 |
| whitestork_02 | -<br>(AE+) | resident | 14,183,321 | 95.6% | 92.8% | 1.6 |
| whitestork_03 | MR07603<br>(AF+) | resident | 21,578,092 | 95.6% | 92.3% | 2.5 |
| whitestork_04 | MR07604<br>(AH+) | resident | 27,452,588 | 95.5% | 92.4% | 3.1 |
| whitestork_05 | MR07605<br>(AJ+) | migrant | 26,009,892 | 95.1% | 92.1% | 2.9 |
| whitestork_06 | MR07606<br>(AK+) | migrant | 16,392,633 | 95.6% | 92.5% | 1.9 |
| whitestork_07 | MR07607<br>(AL+) | resident | 13,998,866 | 94.6% | 91.1% | 1.6 |
| whitestork_08 | MR07608<br>(AM+) | resident | 25,370,010 | 95.2% | 91.9% | 2.9 |
| whitestork_09 | MR07609<br>(AN+) | resident | 14,098,660 | 95.7% | 92.9% | 1.6 |
| whitestork_10 | MR07610<br>(AP+) | resident | 17,334,419 | 95.3% | 92.1% | 2.0 |
| whitestork_11 | MR07611<br>(AR+) | migrant | 9,195,414 | 94.9% | 92.3% | 1.0 |
| whitestork_12 | MR07612<br>(AS+) | resident | 16,196,694 | 95.7% | 92.5% | 1.9 |
| whitestork_13 | MR07613<br>(AT+) | resident | 11,430,485 | 96.0% | 93.2% | 1.3 |
| whitestork_14 | MR07614<br>(AU+) | resident | 22,138,334 | 95.5% | 92.7% | 2.5 |
| whitestork_16 | MR07617<br>(AW+) | resident | 24,063,455 | 94.9% | 92.2% | 2.7 |
| whitestork_17 | MR07616<br>(AY+) | resident | 20,892,640 | 94.9% | 91.7% | 2.4 |
| whitestork_18 | MR9340<br>(E5+) | resident | 18,629,805 | 94.7% | 91.8% | 2.1 |
| whitestork_20 | MR91330<br>(E7+) | resident | 11,178,442 | 94.6% | 92.1% | 1.3 |
| whitestork_21 | MR9134<br>(E8+) | migrant | 27,740,874 | 95.5% | 92.6% | 3.2 |
| whitestork_22 | MR9135<br>(E9+) | resident | 11,331,035 | 94.3% | 91.4% | 1.3 |
| whitestork_23 | MR07602<br>(EA+) | resident | 11,547,520 | 95.3% | 92.2% | 1.3 |
| whitestork_25 | MR09149<br>(EE+) | resident | 23,575,502 | 95.8% | 92.6% | 2.7 |
| whitestork_26 | MR09150<br>(EF+) | resident | 14,152,781 | 95.1% | 92.5% | 1.6 |
| whitestork_27 | -<br>(EJ+) | migrant | 28,865,537 | 95.6% | 92.3% | 3.3 |
| whitestork_28 | -<br>(EK+) | resident | 24,592,060 | 95.0% | 91.9% | 2.8 |
| whitestork_29 | MR07618<br>(EM+) | resident | 18,765,453 | 95.1% | 92.4% | 2.1 |
| whitestork_31 | MR07620<br>(EP+) | resident | 16,893,220 | 95.2% | 92.2% | 1.9 |
| whitestork_32 | MR07621<br>(ER+) | resident | 14,205,130 | 95.6% | 92.8% | 1.6 |
| whitestork_33 | MR07622<br>(ES+) | resident | 29,213,880 | 94.5% | 91.4% | 3.3 |
| whitestork_34 | MR07623<br>(ET+) | resident | 19,338,640 | 95.0% | 92.0% | 2.2 |
| whitestork_35 | MR07630<br>(EU+) | resident | 22,669,186 | 94.9% | 92.3% | 2.6 |
| whitestork_36 | MR07631<br>(EV+) | resident | 22,316,550 | 94.9% | 91.8% | 2.5 |
| whitestork_37 | MR9136<br>(F0+) | resident | 8,412,248 | 93.9% | 91.4% | 0.9 |
| whitestork_39 | MR9138<br>(F2+) | migrant | 24,358,481 | 95.6% | 92.8% | 2.8 |
| whitestork_40 | MR9139<br>(F3+) | resident | 31,316,717 | 95.2% | 92.3% | 3.6 |
| whitestork_41 | MR9140<br>(F4+) | resident | 13,676,717 | 94.2% | 91.7% | 1.5 |
| whitestork_42 | MR9141<br>(F5+) | resident | 11,475,570 | 94.3% | 91.5% | 1.3 |
| whitestork_43 | MR9142<br>(F6+) | resident | 27,164,030 | 95.0% | 91.9% | 3.1 |
| whitestork_44 | MR9143<br>(F7+) | resident | 9,320,085 | 94.9% | 92.4% | 1.1 |
| whitestork_45 | MR9144<br>(F8+) | resident | 33,445,803 | 95.0% | 92.0% | 3.8 |
| whitestork_46 | MR9145<br>(F9+) | resident | 18,048,925 | 94.2% | 91.5% | 2.0 |
| whitestork_48 | MR09088<br>(73+) | resident | 10,073,314 | 95.2% | 92.5% | 1.2 |
| whitestork_49 | MR09089<br>(74+) | migrant | 6,557,210 | 95.1% | 94.4% | 0.8 |
| whitestork_50 | MR09087<br>(8N+) | resident | 20,539,108 | 95.1% | 92.2% | 2.3 |
| whitestork_51 | MR09086<br>(8M+) | resident | 33,004,901 | 95.5% | 92.3% | 3.8 |
| whitestork_52 | MR09090<br>(AA+) | resident | 15,313,648 | 94.6% | 91.7% | 1.7 |
| whitestork_53 | MR09091<br>(AX+) | resident | 17,109,112 | 94.7% | 91.6% | 1.9 |
| whitestork_54 | MR09303<br>(2X+) | migrant | 20,331,304 | 94.0% | 91.3% | 2.3 |
| whitestork_55 | MS03198<br>(K0+) | migrant | 20,444,297 | 95.1% | 92.2% | 2.3 |
| whitestork_56 | MS03199<br>(K1+) | resident | 19,911,560 | 94.8% | 91.7% | 2.2 |
| whitestork_57 | MS03200<br>(K2+) | resident | 8,794,982 | 94.8% | 91.9% | 1.0 |
| whitestork_58 | MS02451<br>(K3+) | resident | 6,704,429 | 94.2% | 92.3% | 0.8 |
| whitestork_60 | MS02453<br>(K5+) | migrant | 22,828,892 | 96.0% | 93.2% | 2.6 |
| whitestork_62 | MS02500<br>(LP+) | migrant | 15,205,512 | 93.6% | 91.2% | 1.7 |

**Table S3.**
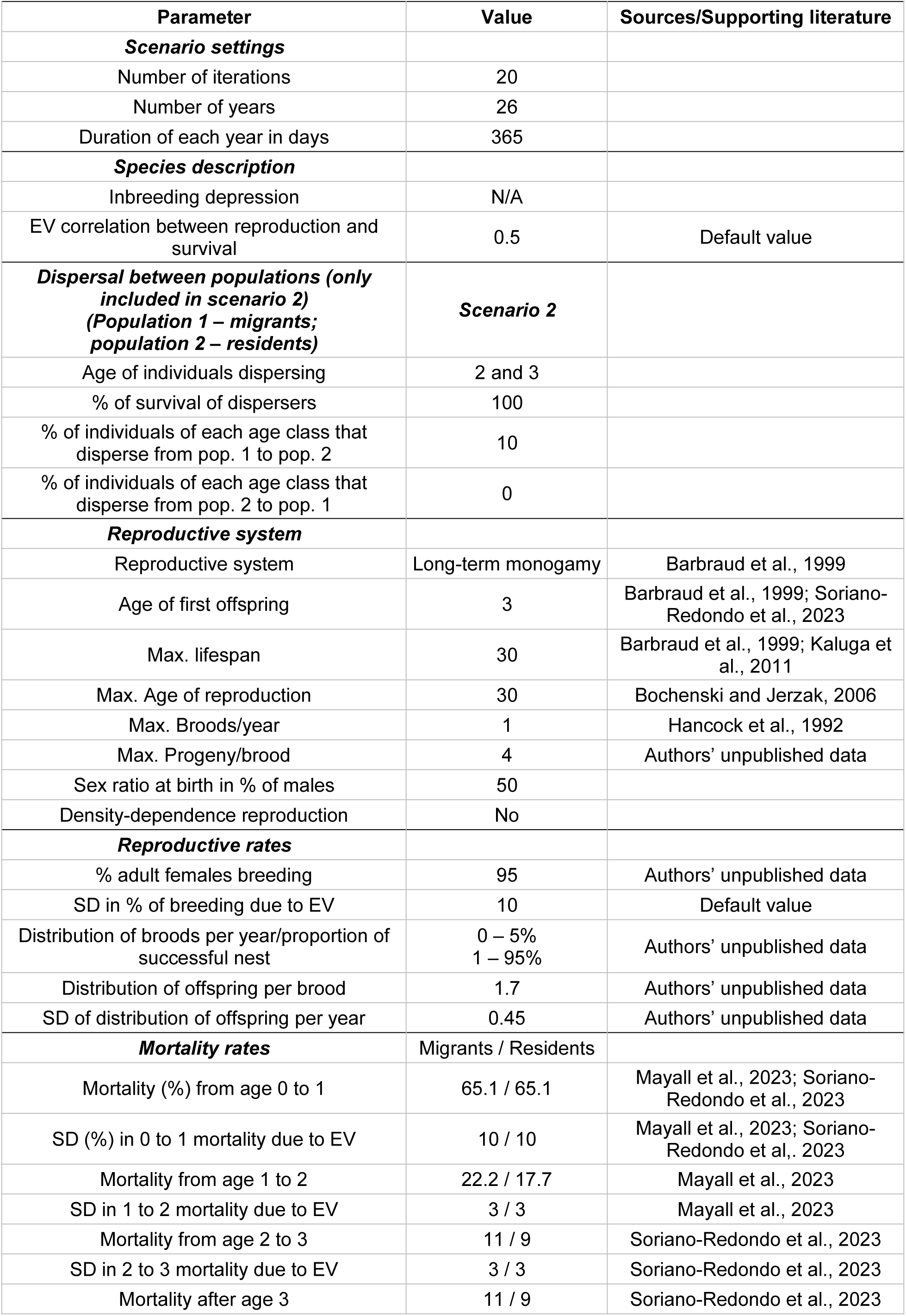
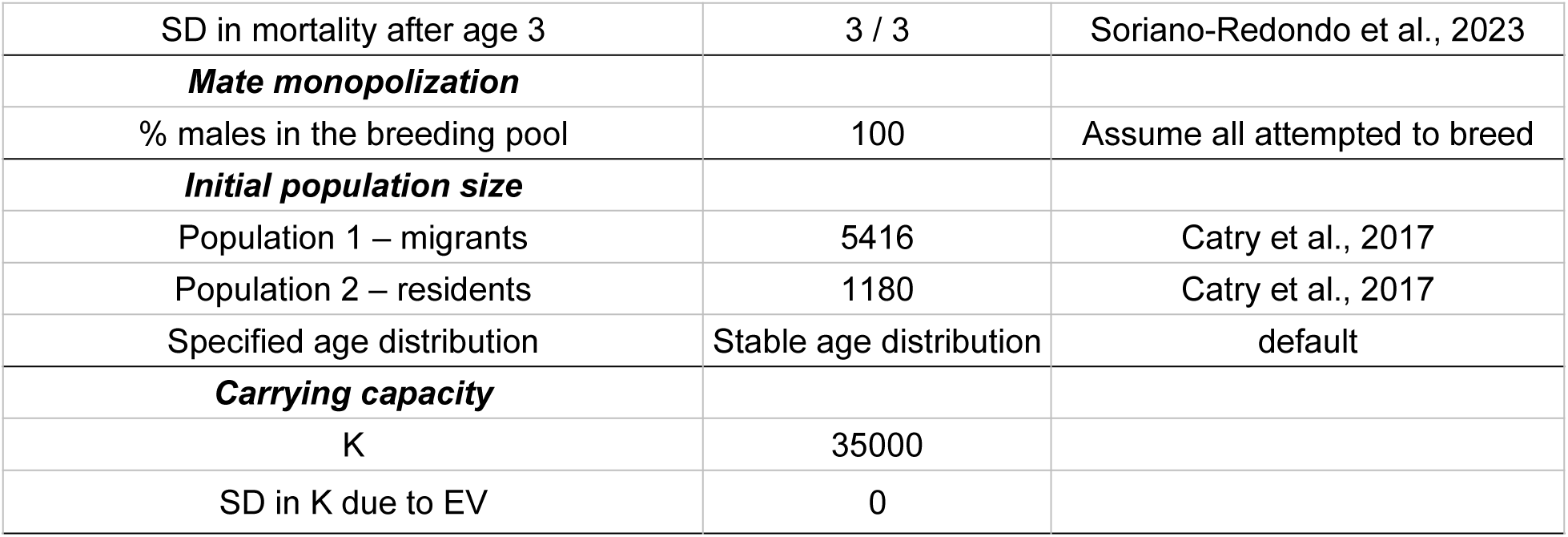
*Vortex10* parameters that were used to model the changes in the number of resident and migratory white storks in Portugal.

**Table S4.** Population demography parameters from *Vortex10* comparing different migratory strategies for the Portuguese white stork population. In scenario 1 individuals do not shift migratory strategy. In scenario 2, 10% of the migratory storks (in the second and third year of life) shift to a resident strategy. Set. r = deterministic growth rate; Stoch. r = stochastic growth rate; SE=standard error; N=population size.

| ***Vortex10* model** | **Description** | **Det. $r$** | **Stoch. $r \pm$**<br>**SE** | **N after 26**<br>**years ± SE** | **% of the**<br>**metapopulation** |
| --- | --- | --- | --- | --- | --- |
| Scenario 1 – no dispersal | Population 1 -<br>migrants | 0.0509 | 0.0506 ±<br>0.0207 | 21087 ±<br>1600 | 73.1 |
| | Population 2 -<br>residents | 0.0746 | 0.0703 ±<br>0.0201 | 7755 ± 550 | 26.9 |
| | Metapopulation | 0.0627 | 0.0556 ±<br>0.0186 | 28842 ±<br>1776 | 100 |
| Scenario 2 – 10% dispersal<br>from the migrant to the<br>resident population | Population 1 -<br>migrants | 0.0509 | 0.0192 ±<br>0.0201 | 9630 ± 859 | 30.2 |
| | Population 2 -<br>residents | 0.0746 | 0.1107 ±<br>0.0199 | 22232 ±<br>1537 | 69.8 |
| | Metapopulation | 0.0627 | 0.0588 ±<br>0.0174 | 31862 ±<br>2024 | 100 |

## REFERENCES

Able, K. P., & Belthoff, J. R. (1998). Rapid ‘evolution’ of migratory behaviour in the introduced house finch of eastern North America. Proceedings of The Royal Society of London. Series B: Biological Sciences, 265(1410), 2063–2071.

Acácio, M., Catry, I., Soriano-Redondo, A., Silva, J. P., Atkinson, P. W., & Franco, A. (2022). Timing is critical: consequences of asynchronous migration for the performance and destination of a long-distance migrant. Movement Ecology, 10(1), 1–16.

Åkesson, S. & Helm, B. (2020). Endogenous programs and flexibility in bird migration. Frontiers in Ecology and Evolution, 8, 78.

Archaux, F., Balança, G., Henry, P. Y., & Zapata, G. (2004). Wintering of white storks in Mediterranean France. Waterbirds, 27(4), 441–445.

Bates, D., Mächler, M., Bolker, B., & Walker, S. (2014). Fitting linear mixed-effects models using lme4. arXiv preprint arXiv:1406.5823.

Bécares, J.; Blas, J.; López-López, P.; Schulz, H.; Torres-Medina, F.; Flack, A.; Enggist, P.; Höfle, U.; Bermejo, A. y, De la Puente, J. (2019). Migración y ecología espacial de la cigüeña blanca en España. Monografía n.° 5 del programa Migra. SEO/BirdLife. Madrid.

Berthold, P. (2001). Ed., Bird Migration: A General Survey (Oxford University Press, Oxford, UK).

Berthold, P., Helbig, A. J., Mohr, G., & Querner, U. (1992). Rapid microevolution of migratory behaviour in a wild bird species. Nature, 360(6405), 668–670.

Both, C., & Visser, M. E. (2001). Adjustment to climate change is constrained by arrival date in a long-distance migrant bird. Nature, 411(6835), 296–298.

Bradnam, K. R., Fass, J. N., Alexandrov, A., Baranay, P., Bechner, M., Birol, I., Boisvert, S., Chapman, J. A., Chapuis, G., Chikhi, R., Chitsaz, H., Chou, W.-C., Corbeil, J., Del Fabbro, C., Docking, T. R., Durbin, R., Earl, D., Emrich, S., Fedotov, P., Fonseca, N. A., et al. (2013). Assemblathon 2: evaluating de novo methods of genome assembly in three vertebrate species. GigaScience, 2(1), 2047–217X.

Byholm, P., Beal, M., Isaksson, N., Lötberg, U., & Åkesson, S. (2022). Paternal transmission of migration knowledge in a long-distance bird migrant. Nature Communications, 13, 1566.

Campioni, L., Dias, M. P., Granadeiro, J. P., & Catry, P. (2020). An ontogenetic perspective on migratory strategy of a long-lived pelagic seabird: Timings and destinations change progressively during maturation. Journal of Animal Ecology, 89(1), 29–43.

Carneiro, C., Gunnarsson, T. G., & Alves, J. A. (2019). Why are whimbrels not advancing their arrival dates into Iceland? Exploring seasonal and sex-specific variation in consistency of individual timing during the annual cycle. Frontiers in Ecology and Evolution, 7, 248.

Catry, I., Encarnação, V., Pacheco, C., Catry, T., Tenreiro, P., da Silva, L. P., & Moreira, F. (2017). Recent changes on migratory behaviour of the White stork (Ciconia ciconia) in Portugal: Towards the end of migration. Airo, 24, 28–35.

Charmantier, A., & Gienapp P. (2014). Climate change and timing of avian breeding and migration: evolutionary versus plastic changes. Evolutionary Applications, 7(1), 15–28.

Charmantier, A., McCleery, R. H., Cole, L. R., Perrins, C., Kruuk, L. E., & Sheldon, B. C. (2008). Adaptive phenotypic plasticity in response to climate change in a wild bird population. Science, 320(5877), 800–803 (2008).

Cheng, Y., Fiedler, W., Wikelski, M., & Flack, A. (2019). “Closer-to-home” strategy benefits juvenile survival in a long-distance migratory bird. Ecology and evolution, 9(16), 8945–8952.

Chernetsov, N., Berthold, P., Querner, U. (2004). Migratory orientation of first-year white storks (*Ciconia ciconia*): inherited information and social interactions. Journal of Experimental Biology, 207(6), 937–943

Conklin, J. R., Lisovski, S., & Battley, P. F. (2021). Advancement in long-distance bird migration through individual plasticity in departure. Nature communications, 12(1), 1–9 (2021).

Curley, S. R., Manne, L. L., & Veit, R. R. (2020). Differential winter and breeding range shifts: Implications for avian migration distances. Diversity and Distributions, 26(4), 415–425.

Dallinga, J. H., & Schoenmakers, S. (1987). Regional decrease in the number of white storks (Ciconia c. ciconia) in relation to food resources. Colonial Waterbirds, 167–177.

DeGiorgio, M., Huber, C. D., Hubisz, M. J., Hellmann, I., & Nielsen, R. (2016). SweepFinder2: increased sensitivity, robustness and flexibility. Bioinformatics, 32(12), 1895–1897.

Delmore, K. E., Toews, D. P., Germain, R. R., Owens, G. L., & Irwin, D. E. (2016). The genetics of seasonal migration and plumage color. Current Biology, 26(16), 2167–2173.

Delmore, K., Illera, J. C., Pérez-Tris, J., Segelbacher, G., Ramos, J. S. L., Durieux, G., Ishigohoka J., & Liedvogel, M. (2020). The evolutionary history and genomics of European blackcap migration. Elife, 9, e54462.

Dias, M. P., Granadeiro, J. P., Phillips, R. A., Alonso, H., Catry, P. (2011). Breaking the routine: individual Cory’s shearwaters shift winter destinations between hemispheres and across ocean basins. Proceedings of the Royal Society B: Biological Sciences, 278(1713), 1786–1793.

Enbody, E. D., Sprehn, C. G., Abzhanov, A., Bi, H., Dobreva, M. P., Osborne, O., Rubin, C.-J., Grant, P. R., Grant, B. R., & Andersson, L. (2021). A multispecies *BCO2* beak color polymorphism in the Darwin’s finch radiation. Current Biology (2021).

Encarnação, V. (2015). “Relatório do VI Censo Nacional de Cegonha-branca *Ciconia ciconia* – 2014” (Tech. Rep. ICNF/CEMPA, Lisboa).

Fay, J. C., & Wu, C. I. (2000). Hitchhiking under positive Darwinian selection. Genetics, 155(3), 1405–1413.

Feng, S., Stiller, J., Deng, Y., Armstrong, J., Fang, Q. I., Reeve, A. H., … & Zhang, G. (2020). Dense sampling of bird diversity increases power of comparative genomics. Nature, 587(7833), 252–257.

Fernández-Cruz, M. (2005). “La migración otoñal de la cigüeña blanca por el Estrecho de Gibraltar” in La Cigüeña Blanca em España. VI Censo Internacional (2004), B. Molina, J. C. Del Moral, Eds. (SEO/BirdLife, Madrid, Spain, pp 162–201).

Ferreira, E., Grilo, F., Mendes, R., Lourenço, R., Santos, S., & Petrucci-Fonseca, F. (2019). Diet of the White Stork (*Ciconia ciconia*) in a heterogeneous Mediterranean landscape: the importance of the invasive Red Swamp Crayfish (*Procambarus clarkii*). Airo, 26, 27–41.

Flack, A., Fiedler, W., Blas, J., Pokrovsky, I., Kaatz, M., Mitropolsky, M., Aghababyan, K., Fakriadis, I., Makrigianni, E., Jerzak, L., Azafzaf, H., Feltrup-Azafzaf, C., Rotics, S., Mokotjomela, T. M., Nathan, R., & Wikelski, M. (2016). Costs of migratory decisions: a comparison across eight white stork populations. Science advances, 2(1), e1500931.

Fraser, K. C., Shave, A., de Greef, E., Siegrist, J., & Garroway, C. J. (2019). Individual variability in migration timing can explain long-term, population-level advances in a songbird. Frontiers in Ecology and Evolution, 7, 324 (2019).

Gilbert, N. I., Correia, R. A., Silva, J. P., Pacheco, C., Catry, I., Atkinson, P. W., Gill, J. A., & Franco, A. M. (2016). Are white storks addicted to junk food? Impacts of landfill use on the movement and behaviour of resident white storks (Ciconia ciconia) from a partially migratory population. Movement Ecology, 4(1), 1–13.

Gill, J. A., Alves, J. A., & Gunnarsson, T. G. (2019). Mechanisms driving phenological and range change in migratory species. Philosophical Transactions of the Royal Society B, 374(1781), 20180047.

Grecian, W. J., Lane, J. V., Michelot, T., Wade, H. M., & Hamer, K. C. (2018). Understanding the ontogeny of foraging behaviour: insights from combining marine predator bio-logging with satellite-derived oceanography in hidden Markov models. Journal of the Royal Society Interface, 15(143), 20180084.

Gu, Z., Pan, S., Lin, Z., Hu, L., Dai, X., Chang, J., Xue, Y., Su, H., Long, J., Sun, M., Ganusevich, S., Sokolov, V., Sokolov, A., Pokrovsky, I., Ji, F., Bruford, M. W., Dixon, A., & Zhan, X. (2021). Climate-driven flyway changes and memory-based long-distance migration. Nature, 1–6.

Hanghøj, K., Moltke, I., Andersen, P. A., Manica, A., & Korneliussen, T. S. (2019). Fast and accurate relatedness estimation from high-throughput sequencing data in the presence of inbreeding. Gigascience, 8(5), giz034.

Hedrick, P. W., & Lacy, R. C. (2015). Measuring relatedness between inbred individuals. Journal of Heredity, 106(1), 20–25.

Hermisson, J., & Pennings, P. S. (2017). Soft sweeps and beyond: understanding the patterns and probabilities of selection footprints under rapid adaptation. Methods in Ecology and Evolution, 8(6), 700–716.

Horton, K. G., La Sorte, F. A., Sheldon, D., Lin, T. Y., Winner, K., Bernstein, G., … & Farnsworth, A. (2020). Phenology of nocturnal avian migration has shifted at the continental scale. Nature Climate Change, 10(1), 63–68.

Horton, K. G., Morris, S. R., Van Doren, B. M., & Covino, K. M. Six decades of North American bird banding records reveal plasticity in migration phenology. Journal of Animal Ecology. doi: 10.1111/1365-2656.13887

Korneliussen, T. S., Albrechtsen, A., & Nielsen, R. (2014). ANGSD: analysis of next generation sequencing data. BMC bioinformatics, 15(1), 1–13.

Levy Karin, E., Mirdita, M., & Söding, J. (2020). MetaEuk—Sensitive, high-throughput gene discovery, and annotation for large-scale eukaryotic metagenomics. Microbiome, 8, 1–15.

Li, H. (2013). Aligning sequence reads, clone sequences and assembly contigs with BWA-MEM. ArXiv:1303.3997 (2013).

Li, H., Handsaker, B., Wysoker, A., Fennell, T., Ruan, J., Homer, N., Marth, G., Abecasis, G., & Durbin, R. (2009). The sequence alignment/map format and SAMtools. Bioinformatics, 25(16), 2078–2079.

Li, H. (2018). Minimap2: pairwise alignment for nucleotide sequences. Bioinformatics, 34(18), 3094–3100.

Liedvogel M. (2019). Genetics of animal and bird migration. In Encyclopedia of Animal Behavior (pp. 323–330). Elsevier (Academic Press).

Lundberg, M., Liedvogel, M., Larson, K., Sigeman, H., Grahn, M., Wright, A., Åkesson, S., & Bensch, S. (2017). Genetic differences between willow warbler migratory phenotypes are few and cluster in large haplotype blocks. Evolution Letters, 1(3), 155–168.

Ma, Y., Ding, X., Qanbari, S., Weigend, S., Zhang, Q., & Simianer, H. (2015). Properties of different selection signature statistics and a new strategy for combining them. Heredity, 115(5), 426–436.

Martins, B. H., Soriano-Redondo, A., Franco, A. M., & Catry, I. (2024). Age mediates access to landfill food resources and foraging proficiency in a long-lived bird species. Animal Behaviour, 207, 23–36.

Mayr, E. (1952). German experiments on orientation of migrating birds. Biological Reviews, 27(4), 394–400.

Meisner, J., & Albrechtsen, A. (2018). Inferring population structure and admixture proportions in low-depth NGS data. Genetics, 210(2), 719–731.

Méndez, V., Gill, J. A., Þórisson, B., Vignisson, S. R., Gunnarsson, T. G., & Alves, J. A. (2021). Paternal effects in the initiation of migratory behaviour in birds. Scientific reports, 11(1), 1–6.

Mueller, T., O’Hara, R. B., Converse, S. J., Urbanek, R. P., & Fagan, W. F. (2013). Social learning of migratory performance. Science, 341(6149), 999–1002.

Nei, M. (1987). Molecular evolutionary genetics. Columbia university press.

Newton, I. (2008). Ed., The migration ecology of birds (Academic Press, London, UK).

Pedersen, L., Jackson, K., Thorup, K., & Tøttrup, A. P. (2018). Full-year tracking suggests endogenous control of migration timing in a long-distance migratory songbird. Behavioral Ecology and Sociobiology, 72(8), 1–10.

Piersma, T., & Drent J. (2003). Phenotypic flexibility and the evolution of organismal design. Trends in Ecology & Evolution, 18(5), 228–233.

Pulido, F., Berthold, P., Mohr, G., & Querner, U. (2001). Heritability of the timing of autumn migration in a natural bird population. Proceedings of the Royal Society of London. Series B: Biological Sciences, 268(1470), 953–959.

Pulido, F. (2007). The genetics and evolution of avian migration. Bioscience, 57(2), 165–174.

Pulido, F., & Berthold, P. (2010). Current selection for lower migratory activity will drive the evolution of residency in a migratory bird population. Proceedings of the National Academy of Sciences, 107(16), 7341–7346.

R Core Team (2020). R: A language and environment for statistical computing. R Foundation for Statistical Computing, Vienna, Austria. URL: https://www.R-project.org/

Rosa, G., Encarnação, V., Pacheco, C. (1998). “Recenseamentos dos efectivos invernantes de Cegonha-branca *Ciconia ciconia* em Portugal (1995-1997)”, in Simpósio sobre Aves Migradoras na Península Ibérica, Costa, L. T., Costa, H., Araújo, M. B., Silva, M. A. Eds. (Sociedade Portuguesa para o Estudo das Aves e Universidade de Évora, Évora, pp 81–85).

Rotics, S., Kaatz, M., Resheff, Y. S., Turjeman, S. F., Zurell, D., Sapir, N., Eggers, U., Flack, A., Fiedler, W., Jeltsch, F., Wikelski, M., & Nathan R. (2016). The challenges of the first migration: movement and behavior of juvenile vs. adult white storks with insights regarding juvenile mortality. Journal of Animal Ecology, 85(4), 938–947.

Rotics, S., Turjeman, S., Kaatz, M., Resheff, Y. S., Zurell, D., Sapir, N., … & Nathan, R. (2017). Wintering in Europe instead of Africa enhances juvenile survival in a long-distance migrant. Animal Behaviour, 126, 79–88.

Rushing, C. S., Royle, J. A., Ziolkowski, D. J., & Pardieck, K. L. (2020). Migratory behavior and winter geography drive differential range shifts of eastern birds in response to recent climate change. Proceedings of the National Academy of Sciences, 117(23), 12897–12903.

Sanchez-Donoso, I., Ravagni, S., Rodríguez-Teijeiro, J. D., Christmas, M. J., Huang, Y., Maldonado-Linares, A., Puigcerver, M., Jiménez-Blasco, I., Andrade, P., Gonçalves, D., Friis, G., Roig, I., Webster, M. T., Leonard, J. A., & Vilà, C. (2022). Massive genome inversion drives coexistence of divergent morphs in common quails. Current Biology, 32(2), 462–469.

Schulz, H., & Schulz, M. (1999). Weißstorch im Aufwind?: Tagungsband internationale Weißstorchtagung, Hamburg, September 26-29, 1996. Bonn: NABU.

Sergio, F., Tanferna, A., Stephanis, De R., Jiménez, L. L., Blas, J., Tavecchia, G., Preatoni, D., & Hiraldo, F. (2014). Individual improvements and selective mortality shape lifelong migratory performance. Nature, 515, 410–413.

Shumate, A., & Salzberg, S. L. (2021). Liftoff: accurate mapping of gene annotations. Bioinformatics, 37(12), 1639–1643.

Simão, F. A., Waterhouse, R. M., Ioannidis, P., Kriventseva, E. V., & Zdobnov, E. M. (2015). BUSCO: assessing genome assembly and annotation completeness with single-copy orthologs. Bioinformatics, 31(19), 3210–3212.

Skotte, L., Korneliussen, T. S., & Albrechtsen, A. (2013). Estimating individual admixture proportions from next generation sequencing data. Genetics, 195(3), 693–702.

Sokolovskis, K., Lundberg, M., Åkesson, S., Willemoes, M., Zhao, T., Caballero-Lopez, V., & Bensch, S. (2023). Migration direction in a songbird explained by two loci. Nature Communications, 14(1), 165.

Soriano-Redondo, A., Acácio, M., Franco, A. M., Herlander Martins, B., Moreira, F., Rogerson, K., & Catry, I. (2020). Testing alternative methods for estimation of bird migration phenology from GPS tracking data. Ibis, 162(2), 581–588.

Soriano-Redondo, A., Franco, A. M., Acácio, M., Martins, B. H., Moreira, F., & Catry, I. (2021). Flying the extra mile pays-off: foraging on anthropogenic waste as a time and energy-saving strategy in a generalist bird. Science of The Total Environment, 146843.

Stanley, C. Q., MacPherson, M., Fraser, K. C., McKinnon, E. A. & Stutchbury, B. J. (2012). Repeat tracking of individual songbirds reveals consistent migration timing but flexibility in route. PloS one, 7(7), e40688

Stoffel, M. A., Nakagawa, S., & Schielzeth, H. (2017). rptR: repeatability estimation and variance decomposition by generalized linear mixed-effects models. Methods in Ecology and Evolution 8, 1639–1644.

Tajima, F. (1989). Statistical method for testing the neutral mutation hypothesis by DNA polymorphism. Genetics, 123(3), 585–595.

Teitelbaum, C. S., Converse, S. J., Fagan, W. F., Böhning-Gaese, K., O’Hara, R. B., Lacy, A. E., & Mueller, T. (2016). Experience drives innovation of new migration patterns of whooping cranes in response to global change. Nature communications, 7(1), 1–7.

Teplitsky, C., Mills, J. A., Alho, J. S., Yarrall, J. W., & Merilä J. (2008). Bergmann’s rule and climate change revisited: Disentangling environmental and genetic responses in a wild bird population. Proceedings of the National Academy of Sciences, 105(36), 13492–13496.

Thorvaldsdóttir, H., Robinson, J. T., & Mesirov, J. P. (2013). Integrative Genomics Viewer (IGV): high-performance genomics data visualization and exploration. Briefings in bioinformatics, 14(2), 178–192.

Toews, D. P., Taylor, S. A., Streby, H. M., Kramer, G. R., & Lovette, I. J. (2019). Selection on VPS13A linked to migration in a songbird. Proceedings of the National Academy of Sciences, 116(37), 18272–18274.

Tortosa, F. S., Caballero, J. M., & Reyes-López, J. (2002). Effect of rubbish dumps on breeding success in the White Stork in southern Spain. Waterbirds, 25(1), 39–43.

Van Vliet, J., Musters, C. J. M., & Ter Keurs, W. J. (2009). Changes in migration behavior of Blackbirds *Turdus merula* from the Netherlands. Bird Study, 56(2), 276–281.

Vardanis, Y., Klaassen, R. H., Strandberg, R., & Alerstam, T. (2011). Individuality in bird migration: routes and timing. Biology letters, 7(4), 502–505 (2011)

Verhoeven, M. A., Loonstra, A. J., Hooijmeijer, J. C., Masero, J. A., Piersma, T., & Senner, N. R. (2018). Generational shift in spring staging site use by a long-distance migratory bird. Biology letters, 14(2), 20170663.

Verhoeven, M. A., Loonstra, A. J., McBride, A. D., Kaspersma, W., Hooijmeijer, J. C., Both, C., Senner, N. R., & Piersma, T. (2021). Age-dependent timing and routes demonstrate developmental plasticity in a long-distance migratory bird. Journal of Animal Ecology, 91(3), 566–579.

Visser, M. E., Perdeck, A. C., Van Balen, J. H., & Both, C. (2009). Climate change leads to decreasing bird migration distances. Global Change Biology, 15(8), 1859–1865.

Wang, M., Zhao, Y., & Zhang, B. (2015). Efficient test and visualization of multi-set intersections. Scientific reports, 5(1), 16923.

Weisenfeld, N. I., Kumar, V., Shah, P., Church, D. M., & Jaffe, D. B. (2017). Direct determination of diploid genome sequences. Genome research, 27(5), 757–767.

## REFERENCES FOR SUPPLEMENTARY MATERIALS

Barbraud, C., Barbraud, J.-C., & Barbraud, M. (1999). Population dynamics of the White Stork *Ciconia ciconia* in western France. Ibis, 141(3), 469–479.

BirdLife International. (2016). Species factsheet: Ciconia ciconia (2015) European Red List Assessment. [Software]. http://datazone.birdlife.org/userfiles/file/Species/erlob/supplementarypdfs/22697691_ciconia_ciconia.pdf

Brook, B. W., Cannon, J. R., Lacy, R. C., Mirande, C., & Frankham, R. (1999, February). Comparison of the population viability analysis packages GAPPS, INMAT, RAMAS and VORTEX for the whooping crane (*Grus americana*). Animal Conservation forum. 2(1), 23–31. 10.1111/j.1469-1795.1999.tb00045.x

Catry, I., Encarnação, V., Pacheco, C., Catry, T., Tenreiro, P., da Silva, L. P., & Moreira, F. (2017). Recent changes on migratory behaviour of the White stork (Ciconia ciconia) in Portugal: towards the end of migration. Airo, 24, 28–35.

Hancock, J., Kushlan, J., & Kahl, P. (1992). Storks, ibises and spoonbills of the world. (London, UK: Academic Press.).

Kalinowski, S. T., & Hedrick, P. W. (1998). An improved method for estimating inbreeding depression in pedigrees. Zoo Biology: Published in affiliation with the American Zoo and Aquarium Association, 17(6), 481–497.

Kanyamibwa, S., Schierer, A., Pradel, R., & Lebreton, J. D. (1990). Changes in adult annual survival rates in a western European population of the White Stork *Ciconia ciconia*. Ibis, 132(1), 27–35.

Lacy, R. C. (2019). Lessons from 30 years of population viability analysis of wildlife populations. Zoo biology, 38(1), 67–77.

Mayall, E., Groves, L., Kennerley, R., Hudson, M., & Franco, A. (2023). Demographic consequences of management actions for the successful reintroduction of the White Stork *Ciconia ciconia* to the UK. Bird Conservation International, 33, e47.

Rotics, S., Kaatz, M., Resheff, Y. S., Turjeman, S. F., Zurell, D., Sapir, N., … & Nathan, R. (2016). The challenges of the first migration: movement and behaviour of juvenile vs. adult white storks with insights regarding juvenile mortality. Journal of Animal Ecology, 85(4), 938–947.

Shephard, J. M., Ogden, R., Tryjanowski, P., Olsson, O., & Galbusera, P. (2013). Is population structure in the European white stork determined by flyway permeability rather than translocation history?. Ecology and Evolution, 3(15), 4881–4895.

Soriano-Redondo, A., Franco, A. M., Acácio, M., Payo-Payo, A., Martins, B. H., Moreira, F., & Catry, I. (2023). Fitness, behavioral, and energetic trade-offs of different migratory strategies in a partially migratory species. Ecology, 104(10), e4151.

Tobolka, M. (2014). Importance of juvenile mortality in Birds’ population: early post-fledging mortality and causes of death in white stork *Ciconia ciconia*. Polish Journal of Ecology, 62(4), 807–813.

Turjeman, S. F., Centeno-Cuadros, A., Eggers, U., Rotics, S., Blas, J., Fiedler, W., … & Nathan, R. (2016). Extra-pair paternity in the socially monogamous white stork (*Ciconia ciconia*) is fairly common and independent of local density. Scientific reports, 6(1), 27976.

